# Determining the probability of hemiplasy in the presence of incomplete lineage sorting and introgression

**DOI:** 10.1101/2020.04.15.043752

**Authors:** Mark S. Hibbins, Matthew J.S. Gibson, Matthew W. Hahn

## Abstract

The incongruence of character states with phylogenetic relationships is often interpreted as evidence of convergent evolution. However, trait evolution along discordant gene trees can also generate these incongruences – a phenomenon known as hemiplasy. Classic phylogenetic comparative methods do not account for discordance, resulting in incorrect inferences about the number of times a trait has evolved, and therefore about convergence. Biological sources of discordance include incomplete lineage sorting (ILS) and introgression, but only ILS has received theoretical consideration in the context of hemiplasy. Here, we derive expectations for the probabilities of hemiplasy and homoplasy with ILS and introgression acting simultaneously. We find that introgression makes hemiplasy more likely than ILS alone, suggesting that methods that account for discordance only due to ILS will be conservative. We also present a method for making statistical inferences about the relative probabilities of hemiplasy and homoplasy in empirical datasets. Our method is implemented in the software package *HeIST* (Hemiplasy Inference Simulation Tool), and estimates the most probable number of transitions among character states given a set of relationships with discordance. *HeIST* can accommodate ILS and introgression simultaneously, and can be applied to phylogenies where the number of taxa makes finding an analytical solution impractical. We apply this tool to two empirical cases of apparent trait convergence in the presence of high levels of discordance, one of which involves introgression between the convergent lineages. In both cases we find that hemiplasy is likely to contribute to the observed trait incongruences.

## Introduction

Convergent evolution of the same phenotype in distantly related species provides some of the most compelling evidence for natural selection. Comparative inferences of convergence require that the species history is known (Felsenstein 1985). Comparative methods applied to such histories implicitly assume that the loci underlying convergent traits also follow the species tree. However, gene trees at individual loci can disagree with each other and with the species tree, a phenomenon known as gene tree discordance. While genomic data allow us to overcome many technical sources of discordance (Delsuc et al. 2005, Dunn et al. 2008, Misof et al. 2014), discordance also has biological causes (Degnan & Rosenberg 2009), and remains a common feature of phylogenomic datasets (Pollard et al. 2006, Fontaine et al. 2015, Pease et al. 2016, Novikova et al. 2016, Wu et al. 2018, Vanderpool et al. 2020).

Gene tree discordance can have multiple sources, including biological causes such as incomplete lineage sorting (ILS), introgression, and horizontal gene transfer, and technical causes such as hidden paralogy or errors in gene tree inference (Schrempf & Szöllősi 2020). Here, we focus primarily on the first two biological causes: ILS and introgression. Looking backwards in time, ILS is the failure of lineages to coalesce within a population before reaching the next most recent ancestral population. The probability of discordance due to ILS is a classic result of the multispecies coalescent, and depends on the population size and the length of time in which coalescence can occur (Hudson 1983, Pamilo & Nei 1988). More recently, the classic multispecies coalescent model has been extended to include introgression (a term we use to encompass hybridization and subsequent gene flow), in a framework called the “multispecies network coalescent” (Yu et al. 2012, Yu et al. 2014, Wen et al. 2016). In this model, species relationships are modelled as a network, with introgression represented by horizontal reticulation edges. Individual loci probabilistically follow or do not follow the reticulation edge, after which they sort according to the multispecies coalescent process (i.e. with ILS). A major advantage of this approach is that ILS and introgression can be modelled simultaneously (reviewed in Degnan 2018), allowing for more detailed study of the consequences of discordance.

Importantly, discordant gene trees can lead to the appearance of apparently convergent traits. This is because discordant gene trees have internal branches that do not exist in the species tree. If a mutation occurs along such a branch at a locus controlling trait variation, it may produce a pattern of character states that is incongruent with the species tree. Incongruent trait patterns are the basis for inferences of convergent evolution (“homoplasy”), and thus this phenomenon has become known as hemiplasy (Avise & Robinson 2008). Since hemiplasy can produce the same kinds of trait incongruence as homoplasy, failing to account for gene tree discordance can generate misleading inferences about convergence (Mendes & Hahn 2016, Mendes et al. 2016). Studies in systems with widespread discordance have found that hemiplasy is a likely explanation for many patterns of incongruence (Copetti et al. 2017, Wu et al. 2018, Guerrero & Hahn 2018).

The problem of hemiplasy makes it clear that robust inferences about the evolution of traits must account for gene tree discordance (Hahn & Nakhleh 2016). Recent work has provided expressions for the probabilities of hemiplasy and homoplasy (Guerrero & Hahn 2018), allowing for an assessment of whether a single transition (hemiplasy) or two transitions (homoplasy) is more likely to explain trait incongruence. This model shows that the most important factors contributing to a high risk of hemiplasy relative to homoplasy are a short internal branch on the species tree (which increases the rate of gene tree discordance), and a low mutation rate (which reduces the probability of the multiple independent transitions needed for homoplasy). However, applying this model in present form to empirical phylogenetic data faces two major limitations. First, incomplete lineage sorting is the only source of gene tree discordance considered, excluding scenarios with gene flow. Second, the model is limited to evolution along a three-taxon tree, restricting calculations for the exact probability of hemiplasy in larger clades.

With genomic data now available for many species, it has become clear that introgression is a common phenomenon (Mallet et al. 2016). Introgression leads to different patterns of gene tree discordance than expected under ILS alone – specifically, minority gene tree topologies supporting a history of introgression are expected to become more common than those produced via ILS alone. These differences form the conceptual basis for common tests of introgression using genomic data (Reich et al. 2009, Green et al. 2010, Durand et al. 2011, Patterson et al. 2012, Pease & Hahn 2015). Introgression also affects the expected coalescence times between pairs of species (Joly et al. 2009, Brandvain et al. 2014, Hibbins & Hahn 2019, Hahn & Hibbins 2019). Pairs of species that have exchanged genes will have lower levels of sequence divergence, and therefore longer shared internal branches, at introgressed loci than expected under ILS alone. These differences in the frequency and branch lengths of genealogies produced by introgression should meaningfully affect the probability of hemiplasy. Therefore, it is important that both sources of gene tree discordance be accounted for in models of trait evolution.

For trees with more than three taxa, the number of possible gene trees and mutational configurations that could explain a particular pattern of trait incongruence increases dramatically. To illustrate this problem, we consider two cases of empirical incongruence of a binary trait. First, consider the case of New Guinea lizards that have evolved green blood from a red-blooded ancestor (Figure 1A; Rodriguez et al. 2018). A clade of 15 taxa contains both the green-blooded species and red-blooded species (the ancestral state). Given the phylogenetic distribution of the six green-blooded species—and no consideration of gene tree discordance—four independent transitions are necessary to explain this incongruence (Figure 1). However, the internal branches on this tree are short and discordance is likely. Individual loci could therefore group the green-blooded taxa into as few as one and as many as six separate clades. Depending on the history at loci affecting blood color, the distribution of green-blooded taxa could therefore be explained by anywhere from one to six mutations, and even more if we consider back-mutations. The one-mutation case represents a single transition due to hemiplasy along a branch that does not exist in the species tree, while the two- and three-mutation cases represent a combination of hemiplasy and homoplasy. The problem becomes even more complex when introgression occurs in the phylogeny, because each reticulation introduces a new set of gene trees formed from the coalescent process at introgressed loci (Hibbins & Hahn 2019). One such example is the origin of a chromosomal inversion spanning a gene involved in wing coloration in the *Heliconius erato/sara* clade of butterflies (Figure 1B; Edelman et al. 2019). Overall, the huge number of possible gene trees (>213 trillion for 15 lizard species; Felsenstein 2004) and the large number of possible mutational events on these trees makes it infeasible to derive an explicit mathematical solution to address questions about hemiplasy in many empirical systems.

**Figure 1:**
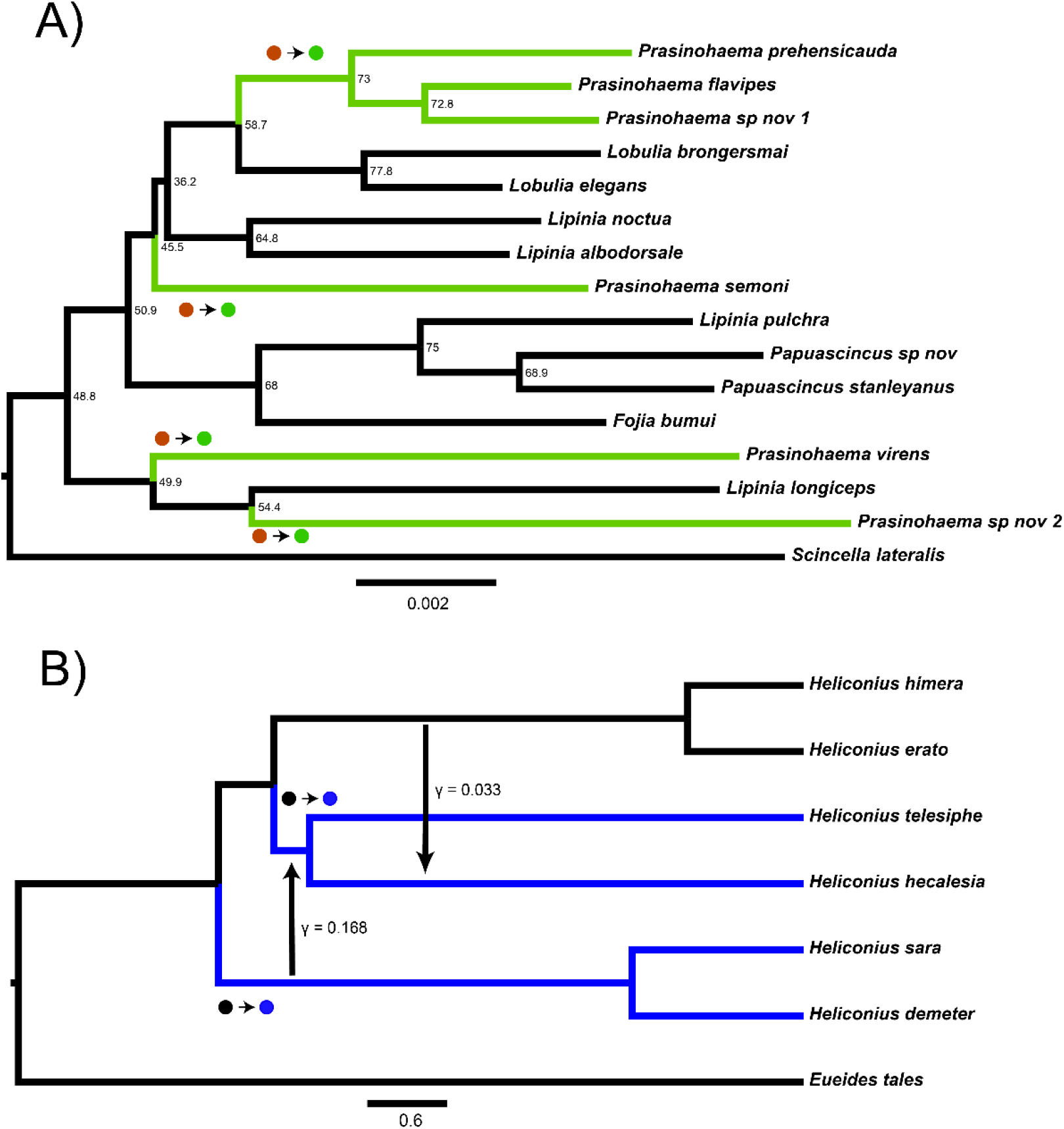
Two empirical examples of apparent convergence in character states that could potentially be explained by hemiplasy. A) Maximum-likelihood species tree of the clade including green-blooded lizards and an outgroup, constructed from the concatenation of 3220 ultra-conserved elements (data from Rodriguez et al. 2018). Branch lengths in substitutions per site; nodes labelled with site concordance factors. B) Coalescent network of *Heliconius erato/sara* clade, processed from the network constructed for the clade in Edelman *et al.* (2019). Branch lengths in units of 2*N* generations; rate, direction, and approximate timing of introgression events indicated by vertical arrows. In both trees, taxa with derived characters are colored, and the most parsimonious transitions from ancestral to derived states are labelled with circles.

Here we make two steps towards addressing these problems. First, we derive expressions for the probabilities of hemiplasy and homoplasy under the multispecies network coalescent for three taxa. Our results show that hemiplasy becomes increasingly likely relative to homoplasy as introgression occurs at a higher rate and at a more recent time relative to speciation. We also show how this pattern is influenced by the direction of introgression. These results highlight the need to account for both sources of discordance in order to understand the origins of a trait incongruence. Second, we present a tool called *HeIST* (<u>He</u>miplasy <u>I</u>nference <u>S</u>imulation <u>T</u>ool) that uses coalescent simulation to dissect patterns of hemiplasy and homoplasy in larger phylogenies. This tool provides an estimate of the most likely number of transitions giving rise to an observed incongruence of binary traits, and accounts for both ILS and introgression. Lastly, we apply *HeIST* to two empirical cases of apparent convergence in a binary trait, finding that hemiplasy is likely to contribute to the observed trait incongruences.

## Materials and Methods

### A model for the probability of hemiplasy under the multispecies network coalescent

To study the effects of introgression on the probability of hemiplasy, we combine concepts from two previously published models: the “parent tree” framework of Hibbins & Hahn (2019), and the model of binary-trait evolution presented in Guerrero & Hahn (2018) (see Wang et al. 2020 for an alternative way to extend the model to incorporate introgression). Consider a rooted three-taxon tree with the topology ((A,B), C). We define *t*_1_ as the time of speciation between lineages A and B in units of 2*N* generations, and *t*_2_ as the time of speciation between C and the ancestor of A and B. We also imagine an instantaneous introgression event between species B and C at time *t*_m_, which can be in either direction (C → B or B → C). We define the total genomic rate of introgression as *δ*, with *δ*_2_ denoting the rate of C → B introgression, and *δ*_3_ the rate of B → C introgression. The history described here is represented by the phylogenetic network shown in Figure 2 (top). Other introgression scenarios can be accommodated by our model (see Discussion), but will not be considered here.

**Figure 2:**
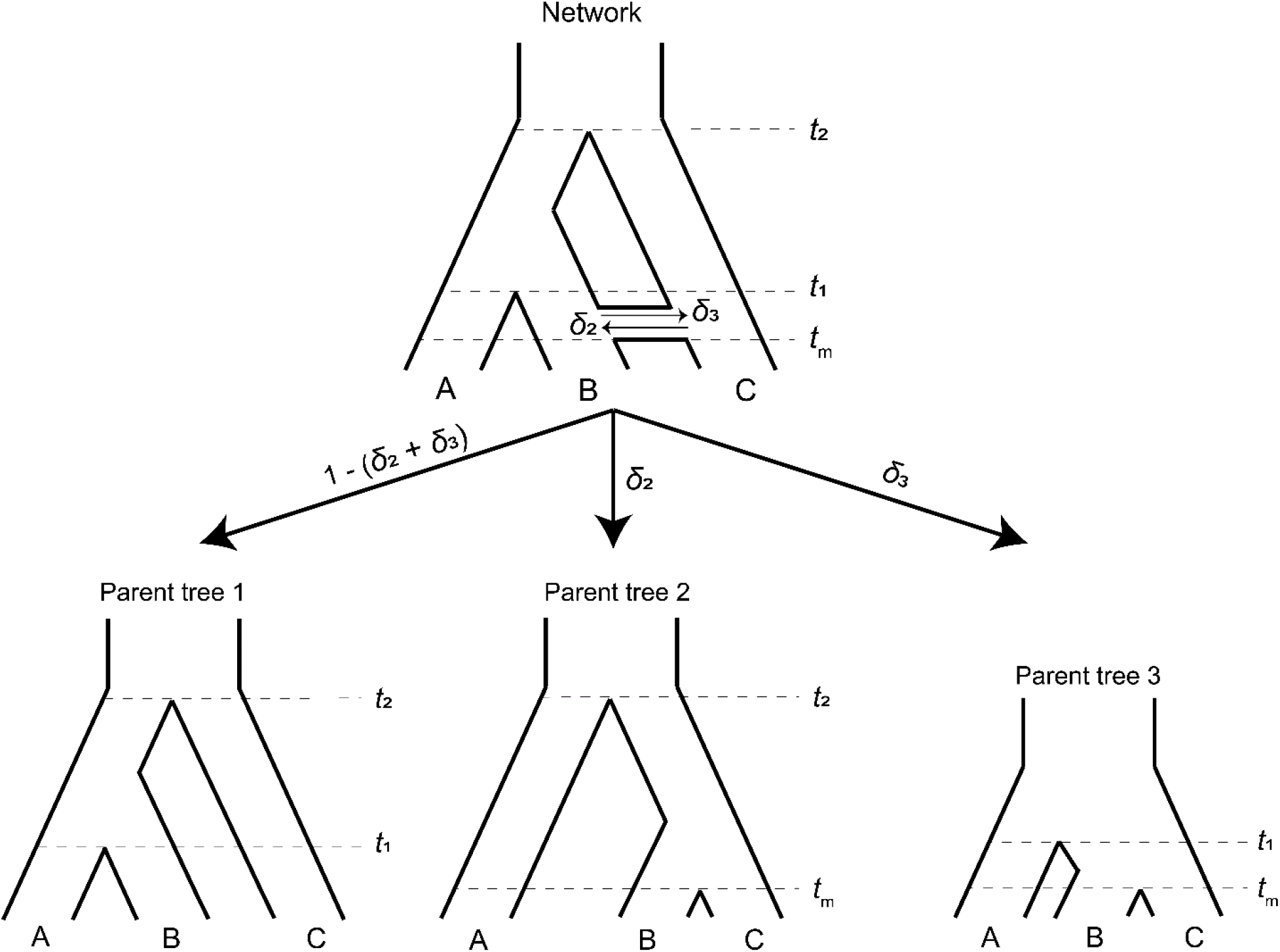
A phylogenetic network (top) can be split into a set of parent trees (bottom) representing the possible histories at individual loci. The probability that a locus is described by a particular parent tree depends on the genomic rate of introgression (arrow labels).

To make it easier to track the history of different gene trees, we imagine that a phylogenetic network can be split into a set of “parent trees” which describe the history at individual loci (Meng & Kubatko 2009, Liu et al. 2014, Hibbins & Hahn 2019) (Figure 2, bottom). Within each of these parent trees, which describe either the species history or the history of introgression, gene trees sort under the multispecies coalescent process. Loci follow the species history, referred to as parent tree 1, with probability 1 − (*δ*_2_ + *δ*_3_). With C → B introgression, some loci will follow the alternative history within parent tree 2, with probability *δ*_2_. In parent tree 2, species B and C are sister and share a “speciation” time of *t*_m_. B → C introgression causes loci to follow parent tree 3 with probability *δ*_3_; in this history lineages B and C are sister and split at time *t*_m_, while the split time of A and the ancestor of B/C is reduced to *t*_1_. This reduction in the second split time in parent tree 3 occurs because the presence of loci from lineage B in lineage C allows C to trace its ancestry through B going back in time. Since B is more closely related to A than C, this allows C to coalesce with A at an earlier time (Figure 2).

Each parent tree can produce four gene trees under the multispecies coalescent process: one tree from lineage sorting, and three equally probable trees from incomplete lineage sorting (Supplementary Figure 1). In other words, introgression always involves ILS, as these are not mutually exclusive histories. Each of these possible gene trees has five branches along which mutations can occur: three tip branches, an internal branch, and an ancestral branch. A subset of these possible gene trees within each parent tree can lead to hemiplasy, while homoplasy can happen in any gene tree (Figure 3). Guerrero & Hahn (2018) provide exact expectations for the probability of a mutation on each branch of each genealogy in an ILS-only model. Before extending this framework to incorporate introgression, the ILS-only model will be briefly described here, using a slightly updated notation that will make it easier to include the effects of introgression.

**Figure 3:**
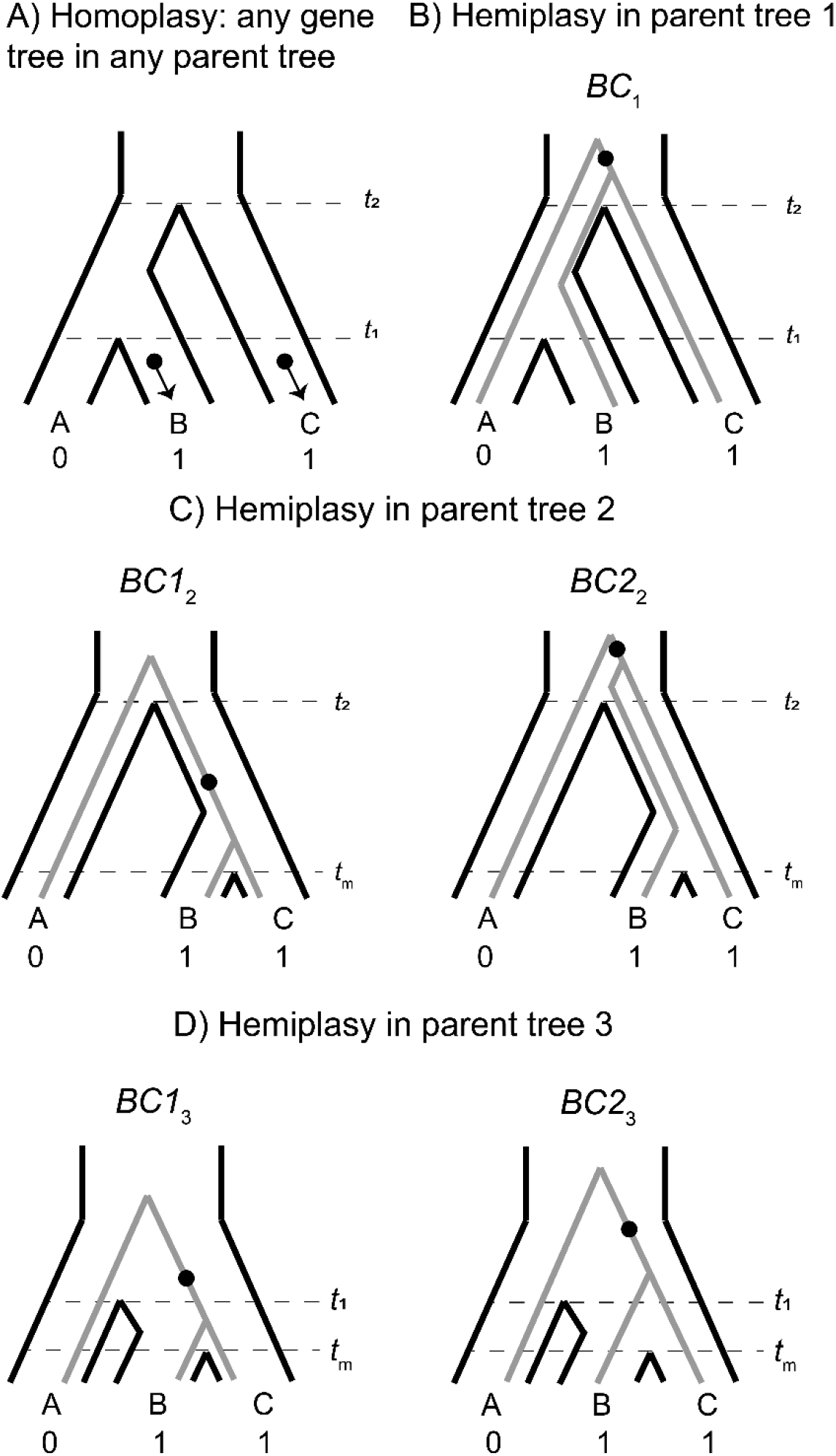
The possible paths to homoplasy and hemiplasy under the multispecies network coalescent. Homoplasy can happen on any gene tree, as long as there are two independent mutations on tip branches (panel A). Homoplasy can also happen via a mutation in the ancestor of all three species, followed by a reversal (not shown). All cases of hemiplasy require a transition on the internal branch of a gene tree with the topology ((B,C),A). In parent tree 1 (panel B), only one such possible gene tree exists (shown in grey; *BC*_1_). In both parent trees 2 and 3 (panels C and D respectively), there are two possible gene trees with this topology. These gene trees differ in internal branch lengths, depending on the parent tree of origin and whether the tree is the result of lineage sorting (*BC*1_2_ and *BC*1_3_) or incomplete lineage sorting (*BC*2_2_ and *BC*2_3_) within introgressed histories.

Consider a binary trait that is incongruent with the described species tree, where species B and C have the derived state and A has the ancestral state. We denote *λ*_1_, *λ*_2_, and *λ*_3_ as the tip branches in any topology leading to species A, B, and C respectively; *λ*_4_ denotes the internal branch of any topology, and *λ*_5_ the branch subtending the root. The notation *ν(λ*, *τ*) represents the probability of a mutation on branch *λ*_i_ in genealogy *τ*, where *τ* represents any of the four gene trees from any of the three parent trees. The rates of 0 → 1 and 1 → 0 mutations are assumed to be equal, and the rate among lineages is assumed to be constant. Finally, to describe individual genealogies, we use the notation *XY*_i=1,2,3_, where *X* and *Y* denote the sister taxa, and the subscript *i* denotes the parent-tree of origin. In cases where a tree topology can be produced by either lineage sorting or ILS, a non-subscripted 1 or 2 is used, respectively. Under the ILS-only model, hemiplasy can only occur through a substitution on branch *λ*_4_ of genealogy *BC*_1_ (Supplementary Figure 1C, Figure 3B). This occurs with the following probability:

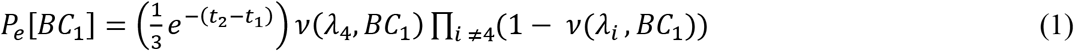

(Guerrero & Hahn, 2018). Equation 1 has three components: the probability of observing genealogy *BC*_1_, the probability that a mutation happens on the internal branch of that genealogy, and the probability that no other mutations occur. See section 1 of the Supplement for the full expressions for each mutation probability.

Now consider the phylogenetic network described earlier and shown in Figure 2. At an introgressed locus, the parent tree topology is ((B,C), A), but could be either parent tree 2 or 3. Within each of these parent trees there are two possible gene trees that share this topology: one produced by lineage sorting (Supplementary Figure 1E, Figure 3C) and one produced by ILS where B and C are still the first to coalesce (Supplementary Figure 1F, Figure 3C). While these trees have the same topology, their expected frequencies and internal branch lengths differ. These quantities also differ depending on the direction of introgression at the locus; i.e. whether the history follows parent tree 2 or 3.

We first consider the C → B direction of introgression, and genealogy *BC*1_2_, which is the result of lineage sorting within parent tree 2. This gives:

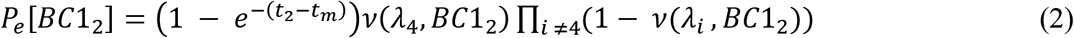

While equation 2 has the same three core components as equation 1, there are several important differences. First, the gene tree probability is the probability of lineage sorting within parent tree 2, which differs from the probability of ILS within parent tree 1. Second, the lower bound of coalescence is *t*_m_ rather than *t*_1_, resulting in a higher probability of lineage sorting in parent tree 2 as compared to parent tree 1. Third, because B and C coalesce more quickly in this tree, they share a longer internal branch, which means the probability of mutation on that branch is higher (see section 1 of the Supplement).

ILS within parent tree 2 produces gene tree *BC*2_2_, in which B and C are the first to coalesce in the common ancestor of all three species. The probability of hemiplasy in this case is:

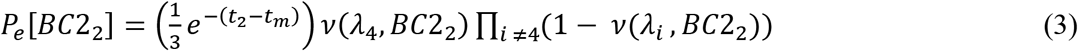

In equation 3, the gene tree probability represents ILS in parent tree 2. This probability is lower than its equivalent in parent tree 1, again because *t*_m_ is the lower bound for coalescence. Since the upper bound to coalescence is the same (*t*_2_), the probability of a mutation on the internal branch of this gene tree is the same as for *BC*_1_ (the ILS topology within parent tree 1). To get the overall probability of hemiplasy due to both ILS and introgression when there is gene flow from C → B, we weight the probability from each gene tree (equations 1–3) by the admixture proportion, giving the following:

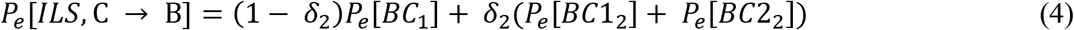

From equation 4, we can see that introgression will increase the probability of hemiplasy over ILS alone (equation 1) whenever the probability of hemiplasy from parent tree 2 is higher than from parent tree 1 (i.e. *P_e_*[*BC*1_2_] + *P_e_*[*BC*2_2_] > *P_e_*[*BC*_1_]). This is true whenever *t*_2_ > *t*_m_ (see section 2 of the Supplement), which is by definition always true in this model.

Finally, we consider the probability of hemiplasy when introgression is in the direction B → C (represented by admixture fraction *δ*_3_). As mentioned previously, this direction of introgression results in an upper bound to coalescence of *t*_1_ rather than *t*_2_. This is the primary difference between the directions of introgression, affecting both the expected gene tree frequencies and mutation probabilities (compare to equations 2 and 3):

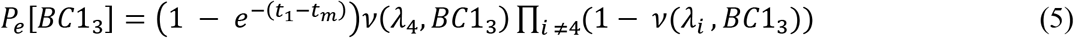

and

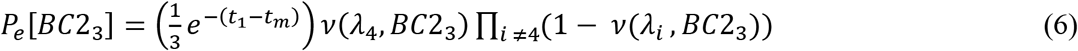

For the general probability of hemiplasy, including both directions of introgression, we now have:

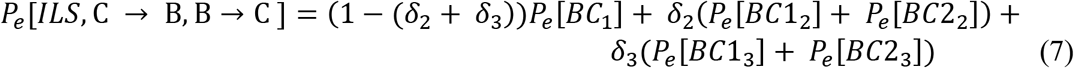

Finally, we consider the probability of homoplasy. As described in Guerrero & Hahn (2018), there are two possible paths to homoplasy for a three-taxon tree where taxa B and C carry the derived state. The first is parallel 0 → 1 mutations on branches *λ*_2_ and *λ*_3_ (Figure 3A), and the second is a 0 → 1 mutation on branch *λ*_5_ followed by a 1 → 0 reversal on branch *λ*_1_. Both of these paths to homoplasy can happen on any possible genealogy, because every topology contains independent tip branches leading to species B and C, as well as an internal branch ancestral to all three species. This gives the following:

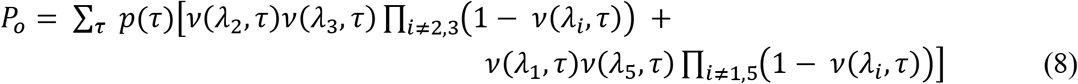

where τ denotes the set of all possible gene trees. (Note that the sum inside equation 8 is multiplied by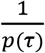 in the main text of Guerrero & Hahn (2018). This is a typo in that paper, but the results presented from their model use the correct expression, *p*(*τ*).) This formulation can also be applied to the extended model with introgression, with the understanding that τ now also includes the gene trees produced by parent trees 2 and 3. Each gene tree used in this summation will have a different set of mutation probabilities, which are detailed in section 1 of the Supplement.

To understand the analytical effect of introgression on the relative risks of hemiplasy and homoplasy, we plotted the ratio *P_e_*/*P_o_* over a realistic range of admixture proportions, timings, and directions (Figure 4). The values of *t*_1_ and *t*_2_ were held constant at 1 and 3.5 coalescent units, respectively, with a population-scaled mutation rate of *θ*= 0.002. These settings ensured a constant contribution of incomplete lineage sorting to the risk of hemiplasy, leading to a baseline ratio of hemiplasy to homoplasy, *P_e_*/*P_o_*, of 0.818 with no introgression. We varied the admixture proportion from 0-10%, and the value of *t*_m_ from 0.99 (just after the most recent speciation) to 0.01, for three different direction conditions: C → B only, B → C only, and equal rates in both directions.

**Figure 4:**
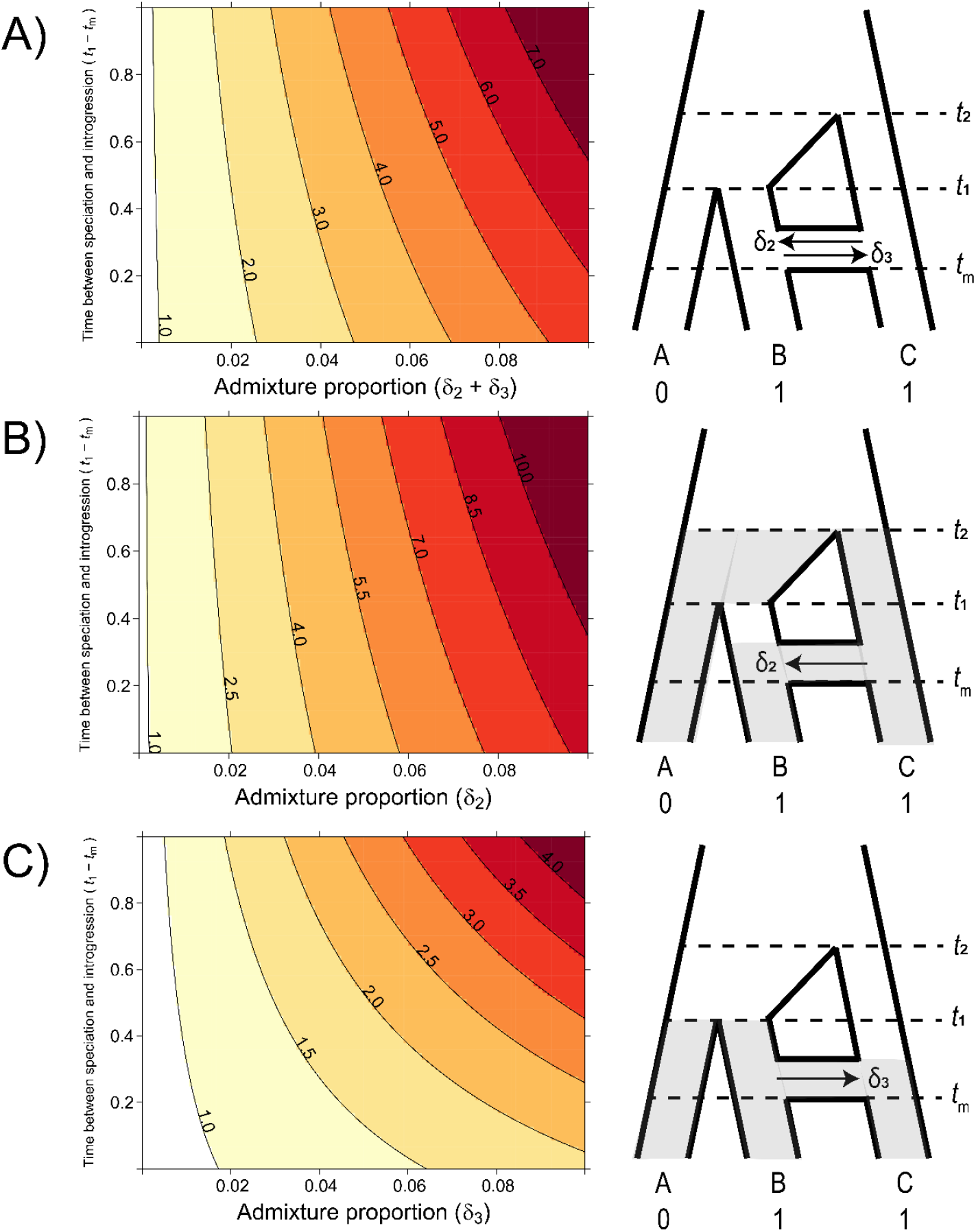
The probability of hemiplasy relative to homoplasy (contours) as a function of the admixture proportion (x-axis), the time between speciation and introgression (y-axis), and the direction of introgression (panels). The contours delineate the factor difference between hemiplasy and homoplasy; for instance, a contour value of 2.0 means hemiplasy is twice as probable as homoplasy in that area of parameter space. At x = 0 in each panel, *P*_e_ / *P*_o_ = 0.818. A) Equal rates of introgression in both directions. B) Introgression in only the C → B direction. Introgression in only the B → C direction.

### *HeIST*: <u>He</u>miplasy <u>I</u>nference <u>S</u>imulation <u>T</u>ool

As described above, it is possible to infer the most likely number of transitions for an incongruent trait while accounting for discordance in a rooted tree with three taxa. However, similar calculations are computationally difficult for larger numbers of taxa. Here, we present a tool built on top of the coalescent simulator *ms* (Hudson 2002) and sequence simulator *Seq-Gen* (Rambaut & Grassly 1997) that provides an intuitive way to interrogate the parameter space of larger trees. Our tool, dubbed *HeIST* (<u>He</u>miplasy <u>I</u>nference <u>S</u>imulation <u>T</u>ool), takes a phylogenetic tree (including an option to specify introgression events) with observed character states as input, and returns a simulated distribution of the number of transitions necessary to explain those character states. Introgression events must be specified as an instantaneous “pulse” from one lineage to another, but we allow flexibility with respect to the timing of that pulse, as well as the rate, direction, and the lineages involved. The input phylogeny must be in coalescent units, but we also include a tool for converting trees given in units of substitutions per site to coalescent units, as long as branches are also associated with concordance factors (see section entitled “Inferring the tip branch lengths of a phylogeny in coalescent units” below).

*HeIST* uses *ms* to simulate a large number of gene trees from the specified species tree or species network, and then simulates the evolution of a single nucleotide site along each of these gene trees using *Seq-Gen*. Loci where the simulated nucleotide states (transformed into 0/1 characters representing ancestral and derived states) match the character states observed on the species tree are taken as replicate simulations of the evolution of the trait being studied. In these “focal” cases, *HeIST* counts the number of mutations that occurred along the gene tree in each simulation. It also returns information on the frequency of tip vs. internal branch mutations, transition vs. reversal mutations, the distribution of gene tree topologies, and whether gene trees originate from the species branching history or introgression history. Finally, it returns a summary of how much hemiplasy is likely to contribute to observed character states, using Fitch parsimony (Fitch 1971) to obtain a homoplasy-only baseline for comparison. *HeIST* is implemented in Python 3 and the package/source code are freely available from https://github.com/mhibbins/HeIST.

### Accuracy of *HeIST*

To confirm that *HeIST* accurately counts mutation events, and is consistent with our theoretical findings, we evaluated its performance under nine simulated conditions with increasing levels of expected hemiplasy. All simulated conditions involve a four-taxon tree with the topology (((4,3),2),1). Species 4 and 2 carry the derived state for a hypothetical binary character. The split of species 1 from the ancestor of 4, 3, and 2 occurs at 8*N* generations in the past. The first three simulated conditions contain no introgression, and progressively decrease the length of the internal branch subtending species 4 and 3. The total tree height was held constant. The simulated internal branch lengths were 2*N*, 1.5*N*, and *N* generations for conditions *ILS1*, *ILS2*, and *ILS3* respectively. The subsequent six conditions maintain the *ILS3* condition for branch lengths, with the addition of an introgression event from species 2 into species 4. For conditions *INT1*, *INT2*, and *INT3*, the timing of introgression was held constant at 0.6*N* generations, while the introgression rate was set to 0.01, 0.05, and 0.1, respectively. For conditions *INT4*, *INT5*, and *INT6*, the introgression rate was held constant at 0.1, while the timing of introgression was reduced to 0.4*N*, 0.2*N*, and 0.1*N*, respectively. The parameters used for each condition are summarized in Supplementary Figure 2.

We performed two sets of simulations: 1) 100 replicates of each condition, consisting of 100,000 gene trees each, with a constant mutation rate of 0.05 per 2*N* generations; 2) 20 replicates of each condition, for each of five different mutation rates per 2*N* generations (0.0005, 0.0025, 0.005, 0.025, 0.05), each consisting of 1,000,000 gene trees. For each combination of parameters, we estimated the probability of hemiplasy conditional on observing the specified trait pattern, and the raw probability of hemiplasy out of the total number of replicates. For the latter simulation set, we estimated the mean-squared error (MSE) using the simulated values as observations and the expected value from theory as the true mean. These MSE values were divided by the simulated mean to compare error across conditions with different ranges of expected values.

### Inferring the tip branch lengths of a phylogeny in coalescent units

Inferences made under the multispecies coalescent require branch lengths specified in coalescent units. However, most standard methods for building phylogenies infer branches in units of substitutions per site. Units of absolute time inferred from substitution rates using molecular clock approaches can be converted into coalescent units, provided that the generation time and effective population size are known. However, these parameters are sometimes not available or accurate for a given system. As an alternative, estimates of gene tree discordance can be used to estimate internal branch lengths in coalescent units, but these provide no information about the lengths of tip branches. For example, the species tree inference software ASTRAL (Zhang et al. 2018b) does not infer tip branch lengths, while the software MP-EST (Liu et al. 2010) adds branches of length 9 for every tip. These tip lengths are necessary to make accurate inferences about hemiplasy and homoplasy from empirical data, since they affect the probability of mutation on tip branches.

To ameliorate this problem, we have applied a simple regression approach for inferring tip lengths in coalescent units (see Bastide et al. 2018 for an alternative method). Our approach makes use of concordance factors: estimates of the fraction of concordant loci with respect to a particular branch in a species tree. Concordance factors come in two flavors: gene concordance factors (gCFs) (Gadagkar et al. 2005, Ane et al. 2007), which estimate the concordance of gene tree topologies, and site concordance factors (sCFs) (Minh et al. 2020a), which do the same for parsimony-informative sites. In general, concordance factors estimated from quartets provide an estimate of 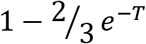, where *T* is the length of the internal branch in coalescent units. With concordance factors given on the internal branches of a tree that has lengths in substitutions per site, the aforementioned formula can be used to obtain estimates of those same branch lengths in units of 2*N* generations. A regression of the internal branch length estimates in both units can then be used to obtain a formula for unit conversion between them. *HeIST* uses this formula to predict the tip branch lengths of the tree in coalescent units. To partially account for uncertainty introduced during tip branch length prediction, *HeIST* can also be run using the lower or upper bounds of the prediction 95% confidence interval as the inferred tip lengths, in addition to the predictions themselves. As a final step in this process, the tree in coalescent units is smoothed, as *ms* requires the input tree to be ultrametric. *HeIST* has two options for how to perform this smoothing. The first redistributes the tree branch lengths so that the distance from the root to each tip is the same; this is done using the convert_to_ultrametric() function in the Python library *ete3* (Huerta-Cepas et al. 2016). The second extends the lengths of tip branches while preserving internal branch lengths; this function is coded within *HeIST*, but was borrowed from a commented block in *ete3*’s source code.

To investigate the potential bias introduced to results from *HeIST* by either phylogenetic inference, the branch regression approach, or subsequent smoothing, we compared the outputs of *HeIST* run from an eight-taxon test tree with known branch lengths. To generate realistic datasets, we first simulated 3000 gene trees from the known species tree using *ms*, and then simulated 1 Kb of sequence from each locus with *θ* = 0.001 using *Seq-Gen*. These loci were concatenated into a single 3 Mb alignment, which was given to RAxML version 8.2.12 (Stamatakis 2014) using the GTR substitution model with rate heterogeneity to infer a species tree in units of substitutions per site. This inferred tree and the concatenated alignment were given to IQ-TREE version 2.0 (Minh et al. 2020b) to infer site concordance factors. This substitution tree with nodes labelled with concordance factors was given as input to *HeIST*, where our branch regression approach was applied. We tested both methods for tree smoothing from this input. The results from this analysis were compared to those obtained using the “true” test tree.

Our regression approach is implemented in *HeIST* and can be run as part of the overall hemiplasy analysis, or separately using the module “*subs2coal*”.

### Empirical applications of *HeIST*

We applied *HeIST* to two empirical case studies where hemiplasy appeared to be a plausible explanation for observed trait incongruences. The first is a dataset of New Guinea lizards (Rodriguez et al. 2018). As described in the Introduction, the genus *Prasinohaema* contains six species that have evolved green blood from a red-blooded ancestor (Figure 1A). Previous analyses of the species tree built from thousands of loci inferred that four independent transitions are necessary to explain the phylogenetic distribution of green-blooded species (Rodriguez et al. 2018). This conclusion is the same using any standard phylogenetic comparative method, whether ancestral state reconstruction is carried out using maximum likelihood (Rodriguez et al. 2018) or Fitch parsimony (this study). However, the phylogeny for this clade contains many short branches (Figure 1), suggesting that a scenario involving at least some hemiplasy (in this case, 1-3 mutations) may be preferred over homoplasy-only scenarios when discordance is accounted for.

To address this question, we used the original dataset of Rodriguez et al. (2018), consisting of 3220 ultra-conserved elements (UCEs) totalling approximately 1.3 Mb for 43 species. We then down-sampled these species to 15 taxa in the clade including the green-blooded species and an outgroup (Figure 1A). We constructed a concatenated maximum likelihood species tree, in addition to gene trees for each UCE, using RAxML version 8.2.12 (Stamatakis 2014). To verify the species tree topology for the 15-taxon subclade, we also constructed a tree with ASTRAL-III version 5.6.3 (Zhang et al. 2018). Site and gene concordance factors were calculated for this tree using IQ-TREE version 2.0 (Minh et al. 2020a, Minh et al. 2020b). To obtain the phylogeny in coalescent units, we employed the regression approach described above for unit conversion as implemented in *HeIST*. The “extend” method was used for tree smoothing. We then used *HeIST* to simulate 10^10^ loci from the lizard subclade containing green-blooded species, with a population-scaled mutation rate (*θ*) of 0.0005 per 2*N* generations. This analysis was performed for each of two outgroups: *Lygosoma sp*, which is sister to 40 species in the 43-species phylogeny, and *Scincella lateralis*, which is sister to the 15-taxon clade containing the green-blooded species. We also calculated *D*-statistics (Green et al. 2010) for 12 trios involving green-blooded taxa, finding no strong evidence of introgression (block bootstrap significance tests, Supplementary Tables 2 and 3). Therefore, our simulations did not include any introgression events.

The second empirical case study involves the origins of a chromosomal inversion spanning a gene important for wing coloration in *Heliconius* butterflies (Edelman et al. 2019). The derived inversion arrangement is shared by four taxa, grouped into two subclades in the *erato/sara* group of *Heliconius*. Fitch parsimony suggests two independent origins, but a combination of short internal branches and introgression between the ancestral populations sharing the inversion (Edelman et al. 2019) strongly suggests a role for hemiplasy. We obtained the phylogenetic network—that is, the species tree with reticulation edges—inferred in units of substitutions per site, in addition to gene concordance factors, from the authors. As our regression approach for conversion to coalescent units cannot be used on phylogenetic networks directly, we used the species tree embedded in the network with concordance factors as input to *subs2coal*. The two most strongly supported introgression events were then added back onto the smoothed network in coalescent units, using the inferred rates and directions, with approximate timings based on the location of the events in the original network and our requirement that these events be instantaneous “pulses.” From this input (shown in Figure 1B), we simulated 10^7^ gene trees in *HeIST* using a mutation rate of 0.0005 per 2*N* generations. We also performed the same simulations without specifying the introgression events to obtain an ILS-only estimate of the probability of hemiplasy.

## Results

### Introgression makes hemiplasy more likely than incomplete lineage sorting alone

Using our model for the probability of hemiplasy and of homoplasy, we examined the ratio *P*_e_ / *P*_o_ over a range of different introgression scenarios. This ratio summarizes how much more probable hemiplasy is than homoplasy for a given area of parameter space; for example, a value of *P*_e_ / *P*_o_ = 2 means hemiplasy is twice as likely as homoplasy. We find that the probability of hemiplasy relative to homoplasy increases as a function of the admixture proportion and how recently introgression occurs relative to speciation (Figure 4). As mentioned in the Introduction, there are several possible reasons for these observed trends. The strongest effect on *P*_e_ / *P*_o_ comes from the admixture proportion: a higher proportion means more loci evolving under parent trees 2 and 3, which means higher frequencies of the genealogies where hemiplasy is possible (i.e. *BC*1_2_, *BC*2_2_, *BC*1_3,_ *BC*2_3_). The range of simulated admixture proportions from 0 to 10 percent was meant to capture a biologically plausible range of values, though rates of introgression can sometimes be much higher than this (e.g. Fontaine et al. 2015). Even in this modest range, the effect on the probability of hemiplasy can be substantial. We found that an admixture proportion of 5% results in hemiplasy being anywhere from 1.5 to 4 times more likely than homoplasy (depending on the timing and direction of introgression; Figure 4). Given the baseline value of *P*_e_/ *P*_o_ with no introgression for our chosen parameters (0.818), this represents at minimum a doubling of the probability of hemiplasy relative to homoplasy.

The effect of the timing of introgression is more complicated, as it manifests in multiple ways. First, more recent introgression increases the values of *t*_2_ − *t*_m_ and *t*_1_ − *t*_m_, which in turn increases the degree of lineage sorting in parent trees 2 and 3, respectively. This leads to a higher frequency of gene trees where hemiplasy is possible. Second, the expected length of the internal branches in these two genealogies increases as introgression becomes more recent, which leads to a higher probability of mutations occurring on these branches. Third, since the total height of each tree is being held constant, more recent introgression reduces the lengths of the tip branches leading to species B and C. This reduces the probability of homoplasy due to parallel substitutions, again making hemiplasy relatively more likely. Finally, the strength of the effect of the timing of introgression increases with the admixture proportion, since it is a property of introgressed loci; in other words, the values of of *t*_2_ − *t*_m_ and *t*_1_ − *t*_m_ do not matter unless loci follow a history of introgression.

The direction of introgression affects the relationship between the admixture proportion, the timing of introgression, and hemiplasy risk (Figures 4B and 4C). While hemiplasy becomes more likely than homoplasy with increased admixture in either direction, *P*_e_ / *P*_o_ is lower in any given part of parameter space for B → C introgression (Figure 4C). This is because the bounds of coalescence for parent tree 3 are *t*_1_ and *t*_m_, which are always closer in time than *t_2_* and *t*_m_ (Figure 2). The smaller internal branch in parent tree 3 leads to a higher rate of ILS, in addition to a shorter internal gene tree branch (and lower mutation probability) on genealogies that undergo lineage sorting in these histories. Finally, the timing of introgression has a stronger effect on *P*_e_ / *P*_o_ in the B → C direction (Figure 4C). This is likely because parent tree 3 is truncated relative to parent tree 2 (see Figure 2), and so the difference *t*_1_ − *t*_m_ makes up a proportionally larger part of the tree height.

### *HeIST* effectively captures the effects of ILS and introgression on hemiplasy risk

To evaluate the performance of *HeIST*, we simulated across nine conditions with increasing expected probabilities of hemiplasy, across five different trait mutation rates. The results, shown in Figure 5, confirm the theoretical predictions shown in Figure 4: the probability of hemiplasy increases as a function of decreasing internal branch length (ILS1-ILS3), increasing rate of introgression (INT1-INT3), and more recent introgression (INT4-INT6). The effect of the timing of introgression is weaker than the effect of the introgression rate, also in line with theoretical expectations. These results held true for both the probability conditional on observing the specified trait pattern (Figure 5A) and the raw probability (Figure 5B).

**Figure 5:**
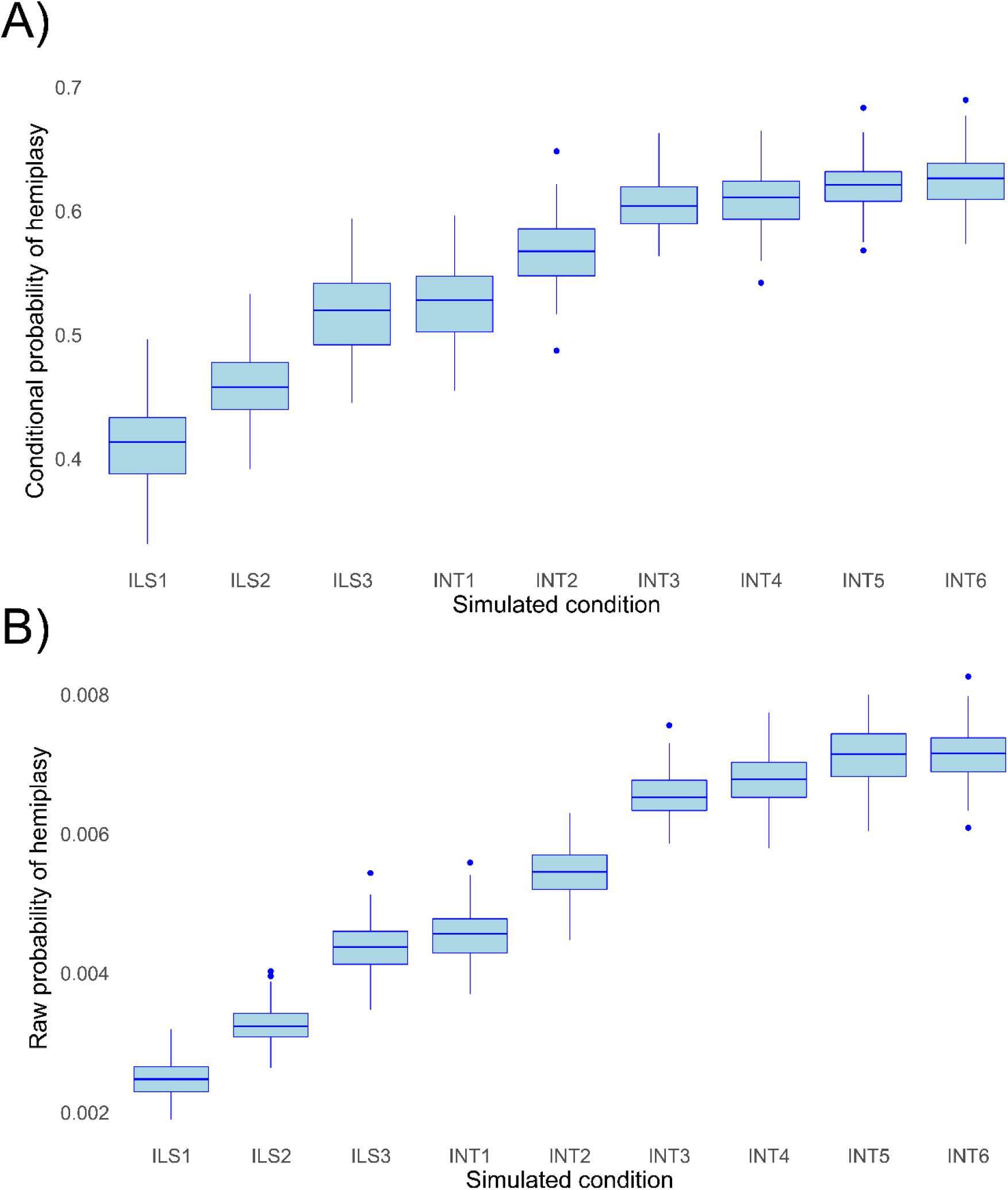
Probabilities of hemiplasy estimated from *HeIST* across nine simulated conditions. ILS1-ILS3 decrease the internal branch length of the species tree; INT1-INT3 introduce introgression between derived taxa with increasing probability; INT4-INT6 make introgression more recent while holding the rate constant. See Supplementary Figure 2 for the exact parameters used in each condition. Panel A shows the probability conditional on observing the trait pattern, while panel B shows the raw probability out of 100,000 simulations.

While the change in the probability of hemiplasy is broadly consistent with theoretical expectations, the probabilities estimated from *HeIST* consistently underestimated the exact values predicted from theory by a small amount (Supplementary Figure 3). We suspect this is due to the occurrence of multiple hits on the same branch of a gene tree, which are not accounted for in our theoretical model. Reversals on branches where hemiplasy can occur would slightly reduce the number of observed hemiplasy cases, leading to the observed underestimation. Consistent with our hypothesis, the mean-normalized mean squared error between simulated and expected values is lower for both lower mutation rates and simulated conditions with a shorter internal branch (Supplementary Figure 4). Overall, the mismatch between simulations and theory appears to be negligible for lower, more realistic trait mutation rates, so we do not believe this will be a concern for most empirical applications.

To evaluate the effects of using *HeIST* on real data, we compared results using the “true” species tree (Supplementary Figure 5A) to those obtained from estimating the species tree with branch lengths using simulated DNA data. This data was run through a pipeline involving estimating a phylogeny using RAxML, converting branch lengths to coalescent units, and smoothing (Supplementary Figure 5B, 5C). In all cases the tree was comprised of eight taxa with no introgression, with three incongruent taxa sharing a hypothetical derived character with a mutation rate of 0.05 per 2*N* generations. Regardless of whether the “extend” or “redistribute” method was used for smoothing, the overall effect of estimating the tree from sequence data was to lengthen both internal and tip branch lengths, reducing the probability of hemiplasy relative to when the true tree was used (see Supplementary Figure 5 for exact probabilities). These results suggest that when our unit-conversion approach and smoothing are applied to empirical datasets, the resulting probability estimates will be conservative with respect to the hypothesis of hemiplasy.

### The distribution of green-blooded New Guinea lizards is likely to have arisen from fewer than four transitions

We investigated the most likely number of transitions to green blood from a red-blooded ancestor in New Guinea lizards of the genus *Prasinohaema* (Rodriguez et al. 2018). Phylogenies constructed using RAxML (Figure 1, Supplementary Figure 6) and ASTRAL (Supplementary Figure 7) recover the phylogeny published by Rodriguez et al. (2018), including the placement of green-blooded species, and also confirm the existence of very short internal branches. In line with this observation, site concordance factors estimated from UCEs indicate very high rates of discordance in this clade, with some approaching a star tree (i.e. all topologies having frequencies of 33%) (Figure 1, Supplementary Table 1). This strongly suggests that the apparent convergent evolution of the green blood phenotype has been affected by hemiplasy.

We used *HeIST* with the 15-taxon subclade containing 6 green-blooded species to determine the most likely number of trait transitions. Using our branch length unit conversion tool *subs2coal*, we obtained a best fit line of *y* = 0.3038 + 157.03*x* with an adjusted *R^2^* of 0.554 (Supplementary Figure 8A). This formula was used to predict the tip branch lengths of the lizard phylogeny in coalescent units, for input to *HeIST* (Supplementary Figure 9). This analysis was repeated using two different outgroups, which differed in their distance from the focal subclade. The results were essentially the same using both outgroups; here we present probabilities using the closer outgroup, *Scincella lateralis*. After simulating 10^10^ loci from the lizard phylogeny using *HeIST*, we obtained 2042 loci with a distribution of derived states that matched the empirical distribution of green-blooded species. It is important to note that this number is expected to be a very small proportion of the total number of simulated loci. This occurs because it is necessary to simulate trait histories randomly, but we use only the ones that match the observed distribution. Due to the enormous space of possibilities, the probability of any single trait distribution will be very low, especially with large numbers of taxa and high rates of trait incongruence.

With four independent transitions required without discordance, there are three possible scenarios that involve at least one hemiplastic transition (Figure 6A). The first is a hemiplasy-only scenario, in which all green-blooded species are grouped into a single monophyletic clade in a gene tree, and a single transition in the ancestor of this clade explains the observed distribution (Figure 6A, left). Out of 2042 focal loci, 726 (35.5%) correspond to this hemiplasy-only case. In the second case, the green-blooded species may be grouped into one or two clades in a gene tree, and there are two independent transitions—at least one of which must involve a discordant ancestral branch (Figure 6A, mid-left). Since there are still multiple independent transitions, this case represents a combination of hemiplasy and homoplasy, but exactly which mutations on which branches are hemiplasy vs. homoplasy will depend on the gene tree topology. 1316 out of 2042 focal cases (64.5%) correspond to this scenario. In the third case, the green-blooded species are grouped into as many as three clades, with three independent transitions, at least one of which must be hemiplastic (Figure 6A, mid-right). Finally, the green-blooded species may be grouped into four clades, with four independent transitions, as in the species tree (Figure 6A, right). We observed no instances of the latter two cases out of 2042 focal loci. These results strongly support the conclusion that, due to hemiplasy, the green-blooded phenotype arose from one or two independent transitions, rather than four.

**Figure 6:**
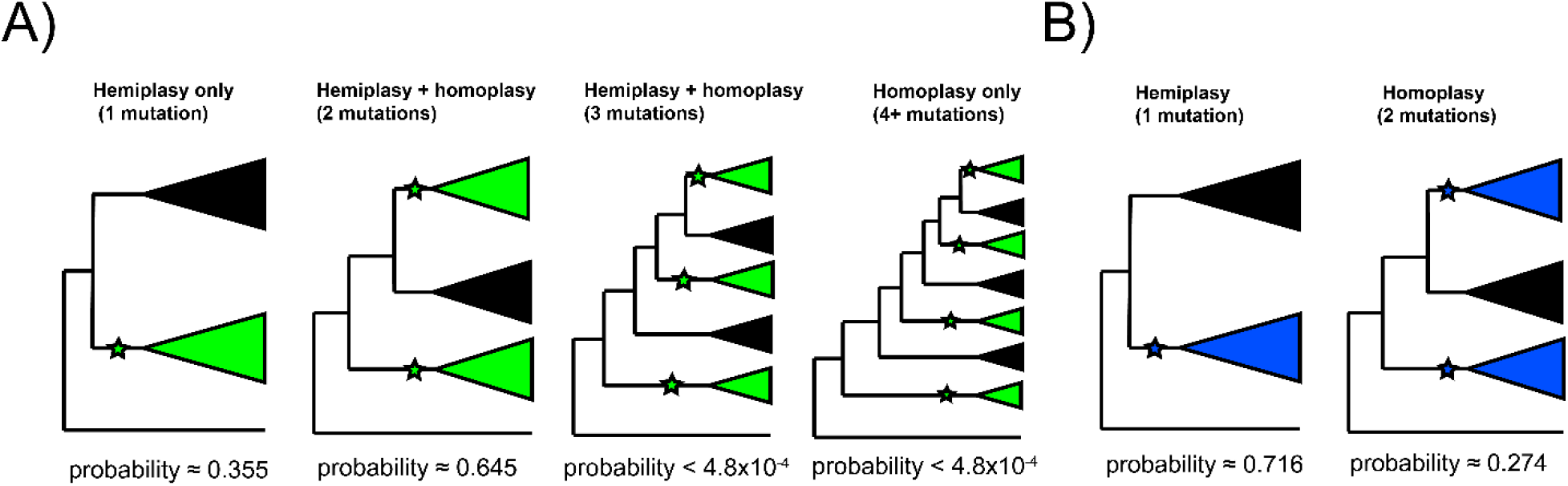
Probable histories for A) the origin of green blood in New Guinea lizards and B) the chromosomal inversion spanning the gene *cortex* in *Heliconius*, calculated using *HeIST*. Trees depict the maximum number of clades expected for gene tree topologies under each scenario, with green-blooded clades in green and inversion clades in blue. Branches with proposed ancestral-to-derived transitions are labelled with stars. Exactly which species are sorted into these clades can vary, meaning many possible gene trees exist for each of the depicted scenarios. Correspondingly, any of the labelled hypothetical mutations could represent hemiplasy or homoplasy (except in the case of a single mutation, which must be hemiplasy), depending on the gene tree topology. Reported probabilities are based on 10^10^ simulated trees for New Guinea lizards and 10^7^ trees for *Heliconius*, with probabilities conditional on matching the empirical trait distributions.

In all 2042 simulated focal cases, the gene trees on which mutations arose grouped the green-blooded species as monophyletic, regardless of the number of mutations that occurred on the tree. In addition, almost all of these monophyletic clades share the same structure, containing two subclades: one containing *P. semoni*, *P. prehensicauda*, *P. flavipes*, and *P. sp nov 1*; another containing *P. virens* and *P. sp nov 2*. It is important to note that the frequency of monophyletic groupings is not expected to reflect the overall distribution of gene trees, but rather the distribution conditional on observing the trait incongruence of interest. These observations make our estimated probabilities easy to interpret: if there was one mutation, it happened in the ancestor of the green-blooded clade; if there were two mutations, they most likely occurred in the ancestors of the two subclades.

Following the logic of “phyloGWAS” (Pease et al. 2016), we checked biallelic sites in the UCE alignment and topologies from the UCE gene trees for a monophyletic clade of green-blooded lizards in order to identify regions potentially associated with variation in blood color. However, both the gene tree and UCE datasets contained missing samples, which made it difficult to confidently identify truly monophyletic clades. On average, approximately 9 taxa were unrepresented in the tips of individual gene trees, and approximately 10 were not assigned a base at individual sites in UCEs. The identity of the missing taxa varied across sites and trees, but often included species inside the 15-taxon subclade containing the green-blooded species, which made it more difficult to consistently polarize and compare patterns of monophyly. In the small proportion of gene trees and UCE sites where information was available for all taxa, we did not find any monophyletic groupings of green-blooded species.

### A chromosomal inversion in the *Heliconius erato*/*sara* clade likely has a single origin

In addition to the analysis of green-blooded lizards, we also investigated the origins of a chromosomal inversion in the *Heliconius erato/sara* clade (Edelman et al. 2019). This inversion spans the gene *cortex*, which is known to influence wing patterning and coloration across butterflies (Joron et al. 2006, Nadeau et al. 2016). While parsimony applied to the species phylogeny would suggest two independent origins of the inversion (Figure 1B), there is clear evidence in Edelman et al. (2019) of both incomplete lineage sorting and introgression among the clades sharing the inversion, implicating a role for hemiplasy.

We inferred branch lengths in coalescent units for the phylogenetic network of these species given in Edelman et al. (2019). Using our unit conversion tool we obtained a best-fit line of *y* = −1.815 + 302.49*x* with an adjusted *R^2^* of 0.98 (though as a note of caution with the *R^2^*, this regression contained only five data points; see Supplementary Figure 8B). The predicted branch leading to the outgroup was extremely long (~ 40*N* generations), so the tree was smoothed using the “extend” method without the outgroup, and the outgroup was re-added post-smoothing at a length proportional to the original network. The two most highly supported introgression events in the inferred phylogenetic network were then added to the coalescent tree with their previously inferred direction, rate, and approximate timing, before being given to *HeIST* as input (Figure 1B).

Using *HeIST*, we found that a single origin of the inversion was most likely, representing 660 of 923 (71.5%) focal cases (Figure 6B, left). The scenario involving two mutations was less likely, but was still found in 253 of 923 cases (27.4%) (Figure 6B, right). We also observed a small number (3/923, 0.32%) of focal cases with three independent transitions. Overall, our results support the original findings of Edelman et al. (2019) that the inversion likely arose once and then was shared between lineages via introgression.

Out of 923 simulated loci matching the trait pattern, we found that 413 originated from an introgressed history. This proportion (0.447) is substantially higher than the sum of introgression probabilities specified in the input (0.201), which suggests that introgression contributes more to the probability of observing the trait incongruence than would be expected by chance. In addition, as in the lizard simulations, we found that almost every simulated focal tree (913/923, 98.9%) grouped the *Heliconius* species that share the inversion as monophyletic. However, there is more variation in the structure of the subclades than there was in the lizards. Nevertheless, we can infer from this that two-mutation cases are most likely to arise as independent mutations in the ancestors of two subclades that are part of a larger monophyletic group.

## Discussion

Phenotypic convergence among species can provide important evidence for natural selection. The molecular variation underlying this convergence can arise through independent mutations at the molecular level (Storz 2016). However, it has recently become clear that such cases of “true” convergence need to be distinguished from cases of apparent convergence due to hemiplasy (Hahn & Nakhleh 2016). Some effort has been made in this regard, through the use of coalescent simulation, summary statistics, and updated comparative approaches (Pease et al. 2016, Copetti et al. 2017, Guerrero & Hahn 2018, Wu et al. 2018). However, these approaches often assume incomplete lineage sorting as the only source of discordance, and cannot explicitly resolve the number of transitions required to explain a trait distribution while accounting for discordance. More recently, Bastide et al. (2018) and Karimi et al. (2020) developed extensions to comparative methods that allow quantitative trait likelihoods to be calculated on phylogenetic networks. However, while phylogenetic network inference methods are often robust to the effects of ILS (Solís-Lemus & Ané 2016, Wen et al. 2018), the estimated networks themselves do not contain the necessary information to simultaneously capture the effects of ILS and introgression on trait probabilities (cf. Mendes et al. 2018).

Here, we take two important steps towards addressing these problems by: 1) studying the effect of introgression on the risk of hemiplasy under the multispecies network coalescent model, and 2) providing a tool that can infer the most probable number of transitions given a phylogenetic distribution of binary traits. We find that introgression increases the risk of hemiplasy over ILS alone, and uncover likely hemiplastic origins for the evolution of green blood from a red-blooded ancestor in New Guinea lizards, and a chromosomal inversion spanning a gene important for wing coloration in *Heliconius*. While our work has important implications for studies of trait evolution, it also carries numerous limitations and simplifying assumptions, which suggest logical next steps for further work. Below we discuss these implications, assumptions, and future directions.

### The probability of hemiplasy due to introgression

A multitude of studies have revealed the potential role of introgression in shaping phenotypic convergence and adaptation (e.g. Heliconius Genome Consortium 2012, Huerta-Sanchez et al. 2014, Jones et al. 2018, Mullen et al. 2020). However, such studies rarely consider how introgression could lead to false inferences of convergence, due to hemiplasy at both the molecular and phenotypic levels, if left unaccounted for. Our model results show that both ILS and introgression must be accounted for in order to make robust inferences of convergent evolution.

Our model for the probability of hemiplasy with introgression, combining concepts from two previously published models, also shares most of their assumptions. First, we have assumed the simplest possible introgression scenario, involving a single pair of species and with introgression occurring instantaneously at some point in the past. However, much more complex introgression scenarios are possible, including introgression between multiple species pairs, involving ancestral populations (and internal branches), at multiple time points in the past, or continuously over a period of time. Horizontal gene transfer, which is more common in prokaryotes, would also require networks that contain reticulation edges spanning very long periods of time. It is not always clear how the probability of hemiplasy would be affected under these alternative introgression scenarios. For example, we assume that the taxa sharing the derived state are also the ones involved in introgression, but introgression between other species pairs could alter patterns of discordance and therefore affect the hemiplasy risk, albeit less directly. Many of these scenarios could be incorporated into the general MSNC framework as additional parent trees, but with more complex histories this may become mathematically intractable even in the three-taxon case; our hemiplasy inference tool, *HeIST*, is designed to ameliorate this issue. Despite these limitations, we can generally expect that introgression will increase the overall risk of hemiplasy whenever rates of introgression are higher between pairs of species that also share the derived state for an incongruent trait. This is because what truly matters is the generation of gene tree topologies with internal branches where hemiplastic transitions can occur; the increased variance in coalescence times under more complex introgression scenarios, while affecting mutation probabilities, should have a comparatively minor effect (Figure 4).

We also assume that the coalescence times and gene tree frequencies of loci underlying trait variation follow neutral expectations, even though alleles controlling trait variation are often under some form of selection. Directional selection on such variation will reduce *N*_e_ relative to neutral expectations, which will decrease the rate of incomplete lineage sorting and consequently hemiplasy due to ILS. Of course, the amount of ILS used in our simulations is not taken directly from neutral expectations, but rather is estimated from real data. Therefore, the effects of selection on traits of interest will only be manifest if they are greater than the general effects of linked selection across the regions used to estimate discordance (cf. Kern & Hahn 2018). On the other hand, introgressed alleles can lead to hemiplasy even in cases where there is no ILS. In fact, directional selection would also make it more likely that introgressed loci have a discordant topology, as it reduces ILS within parent trees 2 and 3. Alternatively, balancing selection can maintain ancestral polymorphism and increase rates of discordance due to ILS. This will also increase the risk of hemiplasy (e.g. Fontaine et al. 2015, Lamichhaney et al. 2016, Palesch et al. 2018).

### Considerations for the inference tool *HeIST*

While the software we introduce here allows for multiple novel types of inferences, it also has several limitations that are important to address. Errors common to all phylogenetic methods can be introduced into the user-specified species tree/network at several steps, including errors in ortholog identification, tree topology, concordance factors, and branch lengths (via both the conversion to coalescent units and tree smoothing). The process of smoothing the coalescent tree should introduce predictable biases in branch length estimates. When using ete3’s method for redistributing branch lengths, internal branches that are very short may have their length increased; conversely, long external branches may be shortened. The lengthening of internal branches decreases the overall rate of discordance, and makes inferences about hemiplasy from *HeIST* conservative. Similarly, when smoothing is done using our function for extending tip branch lengths, the probability of independent mutations on those tip branches (i.e. homoplasy) is increased, again making hemiplasy inferences conservative. The results presented in Supplementary Figure 5 capture the overall effects of errors in phylogeny estimation, branch length prediction, and smoothing.

Errors in inference may affect our approach to branch length unit conversion in several ways. If concordance factors are underestimates—for instance, due to errors in gene tree reconstruction— then the branch lengths in coalescent units will also be underestimates of their true values. The result would be simulations with more ILS and discordance than actually occurred. In cases where there are concerns about branch length estimates, we suggest running *HeIST* across multiple values; for tip branches, the option exists within *HeIST* to use the lower and upper bounds of the prediction interval in addition to the predictions themselves. In addition, if there are tip branch lengths in the original tree that fall outside the range of internal branch length values, the predicted value of those tip branches in coalescent units may be less reliable, since it requires extrapolation beyond the range of datapoints used to fit the regression. Lastly, we note that Bastide et al. (2018) propose an approach to estimating coalescent tip branch lengths on a network using the method of least-squares between pairwise genetic distances and network pairwise distances. We expect this approach to have very similar performance to ours, since linear regression is done using least-squares and pairwise genetic distances should be highly correlated with concordance factors.

There are also several practical points to consider when applying *HeIST* to empirical data. When researchers have questions about hemiplasy involving either very large phylogenies or very low mutation rates, only a small number of simulated trees may match the incongruent pattern found in real data. The large number of simulations required may not be computationally feasible, though careful pruning of species that do not affect inferences of hemiplasy may greatly reduce this limitation. By default, *HeIST* will prune the input phylogeny to include the smallest subclade that contains all the taxa with the derived state, plus a specified outgroup. In addition, while *HeIST* can simulate phylogenies with introgression, it requires that the timing, direction, and rate of each introgression event is provided. To obtain this information, we recommend using a phylogenetic network-based approach such as PhyloNet (Wen et al. 2018), SNaQ (Solís-Lemus & Ané 2016), or the SpeciesNetwork (Zhang et al. 2018a) package within BEAST2 (Bouckaert et al. 2019).

Finally, an issue that concerns both our theoretical work and *HeIST* is the specification of the mutation rate. In both cases, we assume that the rates of 0 → 1 and 1 → 0 transitions are equivalent, and that these rates are constant across the tree under study. Violations of these assumptions will certainly influence the probabilities of hemiplasy and homoplasy, though it is unlikely that underlying mutation rates will vary substantially among closely related lineages (Lynch 2010). More importantly, these rates represent the mutation rate among character states, and may not always be the same as nucleotide mutation rates. We have assumed in the results presented here that transitions between character states are controlled by a single site, and therefore that the nucleotide mutation rate is a good approximation of the trait mutation rate. However, the degree to which this is true will depend on the genetic architecture underlying a trait. For example, transitions in floral color are often underlain by loss-of-function mutations, and many mutational targets can potentially lead to the same phenotypic changes (Rausher 2008, Smith & Rausher 2011). In such cases, the rate of trait transitions can potentially be many times higher than the nucleotide mutation rate, with homoplasy becoming more probable as a result. In contrast, trait transitions can also require multiple molecular changes, the order of which may be constrained by pleiotropy and epistasis. Such changes underlie, for instance, high-altitude adaptation of hemoglobin in mammals (Storz et al. 2009, Tufts et al. 2015). In these cases, the rate of trait transitions may be many times lower than the nucleotide mutation rate, with hemiplasy becoming more probable as a result.

### Evolution of green-blooded lizards & the *Heliconious* inversion

In our analysis of lizards in the genus *Prasinohaema*, we found strong support for one or two independent origins of green blood from a red-blooded ancestor, with two origins being the most likely. This contrasts with analyses that do not account for gene tree discordance, in which four transitions is the best explanation. In *Heliconius*, we found support for a single origin of a chromosomal inversion, in contrast to methods that do not account for discordance. Both of these results strongly suggest that hemiplasy has played a role in the evolution of these traits.

Applications of *HeIST* to these clades involves some system-specific assumptions, the first of which relates to the genetic architecture of the traits under study. For the lizard analysis, it invokes the potentially strong assumption that the green-blooded phenotype is achievable by a single mutation. While the physiological mechanism for this phenotype is well-understood (Austin & Jessing, 1994), the genetic architecture underlying the transition from a red-blooded ancestor is not. As discussed in the previous section, this architecture will affect the choice of *θ* used as the trait evolutionary rate in our simulations. Since the genetic architecture is unknown, our choice of *θ* was based on what is typically observed for nucleotide mutations in vertebrate systems (Lynch 2010). For the *Heliconius* inversion, the architecture is more clear-cut, since chromosomal inversions are a single mutational event by definition. While the per-generation rate of *de novo* chromosomal inversions is not known for many systems, it is certain to be lower than the rate for nucleotide mutations per-site. Nucleotide *θ* is estimated at 0.02-0.03 for *H. melpomene* (Martin et al. 2016), and averages around 0.01 in invertebrates (Lynch 2010). Our choice of *θ* for the inversion was one order of magnitude lower than these estimates.

Another key assumption is that the estimated gene trees and concordance factors are accurate, as is the regression approach for converting branch length units. The observed *R*^2^ of 0.554 for the unit-conversion in the lizard dataset might be interpreted as surprisingly low given that it is a regression of the same quantity measured in two different units. This value likely reflects uncertainty generated in several steps of our analysis, including the estimation of branch lengths in the maximum-likelihood species tree, and the procedure of randomly sampling quartets to estimate sCFs used by IQ-TREE. In *Heliconius*, the *R*^2^ was much higher at 0.98, but with only five data points there was limited information about the true relationship. Nonetheless, we observed the expected positive correlation in both cases, and a sufficient amount of variation is explained to ensure that tip branches estimated in coalescent units are proportionally similar to those in the maximum-likelihood tree, suggesting that the regression approach works well as an approximation. In addition, the regression line on the lizard data appears to slightly over-estimate very short branch lengths in coalescent units, making our inferences of hemiplasy conservative.

## Conclusions

A major question in the study of convergent evolution is whether phenotypic convergence is underlain by convergent changes at the molecular level (Storz 2016). The work presented here is concerned primarily with such molecular changes, and the results of our empirical analyses highlight how apparently convergent phenotypes can arise from a single molecular change. Such shared changes come about as a result of gene tree discordance due to ILS, introgression, or some combination of the two. Given that these phenomena are common in phylogenomic datasets (Pollard et al. 2006, Fontaine et al. 2015, Pease et al. 2016, Novikova et al 2016, Wu et al. 2018), perhaps it should be less surprising that phylogenetically incongruent traits often have a common genetic basis.

Finally, while the tools presented here may help to rule out cases of molecular convergence, the observation of a single molecular origin for a trait does not rule out the occurrence of convergent adaptation in general. Parallel selective pressures from the environment on the same molecular variation may be regarded as one of many possible modes of convergent evolution (Lee & Coop 2017). In studying novel phenotypes such as green blood or wing patterning and coloration, there is still tremendous interest in understanding the ecological pressures that may have led to the independent fixation of single, ancestral changes along multiple lineages. In general, integrative approaches combining modern phylogenomics with an ecological context will pave the way toward an improved understanding of the nature of convergent evolution.

## Acknowledgements

We thank Rafael Guerrero, Leonie Moyle, and Ben Fulton for helpful comments and advice, as well as three anonymous reviewers, the associate editor, and Chris Austin for suggestions that helped improve this work. We also thank Zachary Rodriguez for sharing the lizard data and Nate Edelman for sharing the *Heliconius* phylogenetic network. This work was supported by National Science Foundation grant DEB-1936187.

