## Supplementary Materials for "Determining the probability of hemiplasy in the presence of incomplete lineage sorting and introgression"

Supplementary Materials and Methods

Mark S. Hibbins*, Matthew J.S. Gibson*, and Matthew W. Hahn*†

Department of Biology* and Department of Computer Science†
Indiana University, Bloomington

### 1 Mutation probabilities on genealogies

Each of the twelve possible genealogies under our parent tree model has a set of five branch lengths along which mutations can occur. $\lambda_1$, $\lambda_2$, and $\lambda_3$ denote the tip branches leading to species A, B, and C respectively; $\lambda_4$ denotes the internal branch, and $\lambda_5$ denotes the ancestral branch. As described in the supplement of Guerrero & Hahn (2018), the mutation probability on each of these branches has the general form $\int 1 - e^{-\mu x} f(x)dx$, where $\mu$ is the mutation probability per $2N$ generations, x is the random variable for the branch length, and f(x) is the probability density function for x.

We begin with the mutation probabilities for parent tree 1, which are found in the supplement of Guerrero & Hahn, and will be re-written here to be consistent with notation. In the following notation, parent tree 1 will be denoted as "pt1". Since many of the genealogies are identical in length, the mutation probabilities on their branches can be written with general expressions. We first consider the genealogies $AB2_1$, $BC_1$, and $AC_1$, which are all produced via incomplete lineage sorting in parent tree 1, and share the following set of mutation probabilities:

$$\nu_1[ILS, pt1] = \frac{1}{\Lambda}\int_0^{t_3-t_2}(1 - e^{-\mu(t_1+(t_2-t_1)+x)})\frac{3}{2}(e^{-x} - e^{-3x})dx \tag{1}$$

$$\nu_2[ILS, pt1] = \frac{1}{\Lambda}\int_0^{t_3-t_2}(1 - e^{-\mu(t_1+(t_2-t_1)+x)})3e^{-3x}(1 - e^{-((t_3-t_2)-x)})dx \tag{2}$$

$$\nu_4[ILS, pt1] = \frac{1}{\Lambda}\int_0^{t_3-t_2}3e^{-3y}(\int_0^{(t_3-t_2)-y}e^{-x}(1 - e^{-\mu x})dx)dy \tag{3}$$

$$\nu_5[ILS, pt1] = \frac{1}{\Lambda}\int_0^{t_3-t_2}(1 - e^{-\mu((t_3-t_2)-x)})3e^{-3x}dx \tag{4}$$

In each of the above, $\Lambda = 1 + \frac{1}{2}e^{-3(t_3-t_2)} - \frac{3}{2}e^{-(t_3-t_2)}$ is the probability of coalescence of A, B, and C in their ancestral population. $t_3$ denotes the total height of the tree, i.e. the time at the base of the tree. The difference between $t_3$ and $t_2$ determines the duration of the ancestral population of all three taxa, before speciation occurs. Equations 1 through 4 each represent the mutation probabilities for multiple branches, which are as follows:

$$\nu_1[ILS, pt1] = \nu(\lambda_3, AB2_1) = \nu(\lambda_1, BC_1) = \nu(\lambda_2, AC_1) \tag{5}$$

$$\nu_2[ILS, pt1] = \nu(\lambda_1, AB2_1) = \nu(\lambda_2, AB2_1) = \nu(\lambda_2, BC_1) = \nu(\lambda_3, BC_1) = \nu(\lambda_1, AC_1) = \nu(\lambda_3, AC_1) \quad (6)$$

$$\nu_4[ILS, pt1] = \nu(\lambda_4, AB2_1) = \nu(\lambda_4, BC_1) = \nu(\lambda_4, AC_1) \quad (7)$$

$$\nu_5[ILS, pt1] = \nu(\lambda_5, AB2_1) = \nu(\lambda_5, BC_1) = \nu(\lambda_5, AC_1) \quad (8)$$

The gene tree produced by lineage sorting in parent tree 1, $AB1_1$, has a different set of mutation probabilities, since the branches have different expected lengths. These are:

$$\nu(\lambda_1, AB1_1) = \nu(\lambda_2, AB1_1) = \frac{1}{1 - e^{-(t_2 - t_1)}} \int_0^{t_2 - t_1} (1 - e^{-\mu(t_1 + x)}) e^{-x} dx \quad (9)$$

$$\nu(\lambda_3, AB1_1) = \frac{1}{1 - e^{-(t_3 - t_2)}} \int_0^{t_3 - t_2} (1 - e^{-\mu(t_1 + (t_2 - t_1) + x)}) e^{-x} dx \quad (10)$$

$$\nu(\lambda_4, AB1_1) = \int_0^{t_2 - t_1} \frac{e^{-y}}{1 - e^{-(t_2 - t_1)}} (\int_0^{t_3 - t_2} (1 - e^{-\mu((t_2 - t_1) - y + x)}) \frac{e^{-x}}{1 - e^{-(t_3 - t_2)}} dx) dy \quad (11)$$

$$\nu(\lambda_5, AB1_1) = \frac{1}{1 - e^{-(t_3 - t_2)}} \int_0^{t_3 - t_2} (1 - e^{-\mu((t_3 - t_2) - x)}) e^{-x} dx \quad (12)$$

Now we consider introgression, starting with parent tree 2. Many of the mutation probabilities are symmetrical with parent tree 1 and therefore remain the same, and the remainder have the same general form with different parameters. For the ILS genealogies $BC2_2$, $AB_2$, and $AC_2$, equations 1 and 2 have the time of A-B speciation ($t_1$) replaced with the timing of B-C introgression ($t_m$). This gives:

$$\nu_1[ILS, pt2] = \frac{1}{\Lambda} \int_0^{t_3 - t_2} (1 - e^{-\mu(t_m + (t_2 - t_m) + x)}) \frac{3}{2} (e^{-x} - e^{-3x}) dx \quad (13)$$

$$\nu_2[ILS, pt2] = \frac{1}{\Lambda} \int_0^{t_3 - t_2} (1 - e^{-\mu(t_m + (t_2 - t_m) + x)}) 3e^{-3x} (1 - e^{-((t_3 - t_2) - x)}) dx \quad (14)$$

$$\nu_4[ILS, pt2] = \nu_4[ILS, pt1] \quad (15)$$

$$\nu_5[ILS, pt2] = \nu_5[ILS, pt1] \quad (16)$$

These correspond to the following branch mutation probabilities:

$$\nu_1[ILS, pt2] = \nu(\lambda_1, BC2_2) = \nu(\lambda_3, AB_2) = \nu(\lambda_2, AC_2) \quad (17)$$

$$\nu_2[ILS, pt2] = \nu(\lambda_2, BC2_2) = \nu(\lambda_3, BC2_2) = \nu(\lambda_1, AB_2) = \nu(\lambda_2, AB_2) = \nu(\lambda_1, AC_2) = \nu(\lambda_3, AC_2) \quad (18)$$

$$\nu_4[ILS, pt2] = \nu(\lambda_4, BC2_2) = \nu(\lambda_4, AB_2) = \nu(\lambda_4, AC_2) \quad (19)$$

$$\nu_5[ILS, pt2] = \nu(\lambda_5, BC2_2) = \nu(\lambda_5, AB_2) = \nu(\lambda_5, AC_2) \quad (20)$$

For the genealogy produced by lineage sorting in parent tree 2, $BC1_2$, we have:

$$\nu(\lambda_2, BC1_2) = \nu(\lambda_3, BC1_2) = \frac{1}{1 - e^{-(t_2 - t_m)}} \int_0^{t_2 - t_m} (1 - e^{-\mu(t_m + x)}) e^{-x} dx \tag{21}$$

$$\nu(\lambda_1, BC1_2) = \frac{1}{1 - e^{-(t_3 - t_2)}} \int_0^{t_3 - t_2} (1 - e^{-\mu(t_m + (t_2 - t_m) + x)}) e^{-x} dx \tag{22}$$

$$\nu(\lambda_4, BC1_2) = \int_0^{t_2 - t_m} \frac{e^{-y}}{1 - e^{-(t_2 - t_m)}} (\int_0^{t_3 - t_2} (1 - e^{-\mu((t_2 - t_m) - y + x)}) \frac{e^{-x}}{1 - e^{-(t_3 - t_2)}} dx) dy \tag{23}$$

$$\nu(\lambda_5, BC1_2) = \nu(\lambda_5, AB1_1) \tag{24}$$

Finally, we consider parent tree 3. The mutation probabilities have the same formulation as parent tree 2, with two key changes: since parent tree 3 is shorter (Figure 2 of main text), $t_2$ is replaced by $t_1$. This also applies to the value of $\Lambda$, which we will denote for parent tree 3 as $\Lambda_3 = 1 + \frac{1}{2}e^{-3(t_3 - t_1)} - \frac{3}{2}e^{-(t_3 - t_1)}$. For the ILS genealogies $BC2_3$, $AB_3$, and $AC_3$, this gives:

$$\nu_1[ILS, pt3] = \frac{1}{\Lambda_3} \int_0^{t_3 - t_1} (1 - e^{-\mu(t_m + (t_1 - t_m) + x)}) \frac{3}{2} (e^{-x} - e^{-3x}) dx \tag{25}$$

$$\nu_2[ILS, pt3] = \frac{1}{\Lambda_3} \int_0^{t_3 - t_1} (1 - e^{-\mu(t_m + (t_1 - t_m) + x)}) 3e^{-3x} (1 - e^{-((t_3 - t_1) - x)}) dx \tag{26}$$

$$\nu_4[ILS, pt3] = \frac{1}{\Lambda_3} \int_0^{t_3 - t_1} 3e^{-3y} (\int_0^{(t_3 - t_1) - y} e^{-x} (1 - e^{-\mu x}) dx) dy \tag{27}$$

$$\nu_5[ILS, pt3] = \frac{1}{\Lambda_3} \int_0^{t_3 - t_1} (1 - e^{-\mu((t_3 - t_1) - x)}) 3e^{-3x} dx \tag{28}$$

Where:

$$\nu_1[ILS, pt3] = \nu(\lambda_1, BC2_3) = \nu(\lambda_3, AB_3) = \nu(\lambda_2, BC_3) \tag{29}$$

$$\nu_2[ILS, pt3] = \nu(\lambda_2, BC2_3) = \nu(\lambda_3, BC2_3) = \nu(\lambda_1, AB_3) = \nu(\lambda_2, AB_3) = \nu(\lambda_1, AC_3) = \nu(\lambda_3, AC_3) \tag{30}$$

$$\nu_4[ILS, pt3] = \nu(\lambda_4, BC2_3) = \nu(\lambda_4, AB_3) = \nu(\lambda_4, AC_3) \tag{31}$$

$$\nu_5[ILS, pt3] = \nu(\lambda_5, BC2_3) = \nu(\lambda_5, AB_3) = \nu(\lambda_5, AC_3) \tag{32}$$

Finally, for the genealogy $BC1_3$, the mutation probabilities are as follows:

$$\nu(\lambda_2, BC1_3) = \nu(\lambda_3, BC1_3) = \frac{1}{1 - e^{-(t_1 - t_m)}} \int_0^{t_1 - t_m} (1 - e^{-\mu(t_m + x)}) e^{-x} dx \tag{33}$$

$$\nu(\lambda_1, BC1_3) = \frac{1}{1 - e^{-(t_3 - t_1)}} \int_0^{t_3 - t_1} (1 - e^{-\mu(t_m + (t_1 - t_m) + x)}) e^{-x} dx \tag{34}$$

$$\nu(\lambda_4, BC1_3) = \int_0^{t_1 - t_m} \frac{e^{-y}}{1 - e^{-(t_1 - t_m)}} (\int_0^{t_3 - t_1} (1 - e^{-\mu((t_1 - t_m) - y + x)}) \frac{e^{-x}}{1 - e^{-(t_3 - t_1)}} dx) dy \tag{35}$$

$$\nu(\lambda_5, BC1_3) = \frac{1}{1 - e^{-(t_3 - t_1)}} \int_0^{t_3 - t_1} (1 - e^{-\mu((t_3 - t_1) - x)}) e^{-x} dx \tag{36}$$

#### 2 When does introgression makes hemiplasy more likely than ILS alone?

The probability of hemiplasy with $C \to B$ introgression is

$$P_e = (1-\delta)P_e[BC_1] + \delta(P_e[BC1_2] + P_e[BC2_2]) \tag{37}$$

From this, it can be seen that introgression makes hemiplasy more likely than ILS alone when:

$$P_e[BC1_2] + P_e[BC2_2] > P_e[BC_1] \tag{38}$$

When is this true? Substituting the relevant expressions from the main text gives:

$$\begin{aligned}(1-e^{-(t_2-t_m)})v(\lambda_4, BC1_2)\prod_{i\neq 4}(1-v(\lambda_i, BC1_2))+ \\ (\frac{1}{3}e^{-(t_2-t_m)})v(\lambda_4, BC2_2)\prod_{i\neq 4}(1-v(\lambda_i, BC2_2)) \\ > (\frac{1}{3}e^{-(t_2-t_1)})v(\lambda_4, BC_1)\prod_{i\neq 4}(1-v(\lambda_i, BC_1))\end{aligned} \tag{39}$$

$$\begin{aligned}(1-e^{-(t_2-t_m)})v(\lambda_4, BC1_2)\prod_{i\neq 4}(1-v(\lambda_i, BC1_2)) > \\ (\frac{1}{3}e^{-(t_2-t_1)})v(\lambda_4, BC_1)\prod_{i\neq 4}(1-v(\lambda_i, BC_1))- \\ (\frac{1}{3}e^{-(t_2-t_m)})v(\lambda_4, BC2_2)\prod_{i\neq 4}(1-v(\lambda_i, BC2_2))\end{aligned} \tag{40}$$

The mutation probabilities on the right side of the inequality are equal since they are on the same topology with the same branch lengths. Therefore, equation 40 can simplified:

$$\begin{aligned}(1-e^{-(t_2-t_m)})v(\lambda_4, BC1_2)\prod_{i\neq 4}(1-v(\lambda_i, BC1_2)) > \\ v(\lambda_4, BC_1)\prod_{i\neq 4}(1-v(\lambda_i, BC_1))(\frac{1}{3}e^{-(t_2-t_1)} - \frac{1}{3}e^{-(t_2-t_m)})\end{aligned} \tag{41}$$

As the most conservative case, let us assume a hybrid speciation scenario where $t_1 = t_m$. This represents the most conservative introgression scenario, since Figure 4 in the main text shows that more recent introgression makes hemiplasy more likely. In this case, the right side of the inequality simplifies to 0, leaving

$$(1-e^{-(t_2-t_m)})v(\lambda_4, BC1_2)\prod_{i\neq 4}(1-v(\lambda_i, BC1_2)) > 0 \tag{42}$$

This is true whenever $t_2 > t_m$, which is true by definition in this model. Therefore, introgression always makes hemiplasy more likely than ILS alone.

#### 3 Supplementary Figures and Tables

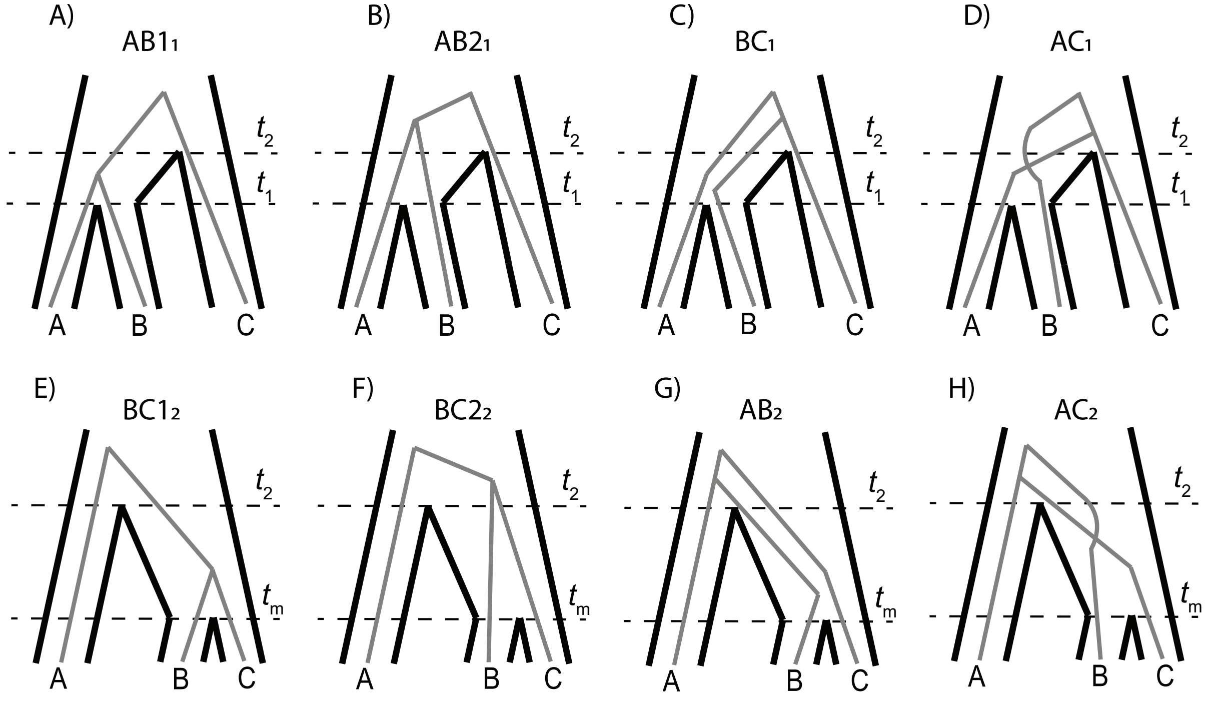


Supplementary Figure 1: Each parent tree in our model generates four gene trees; one generated from lineage sorting (Panels A and E), and three equally likely trees generated from incomplete lineage sorting (panels B-D, F-H). Top row shows the trees generated from parent tree 1; the bottom row shows the trees for parent tree 2.

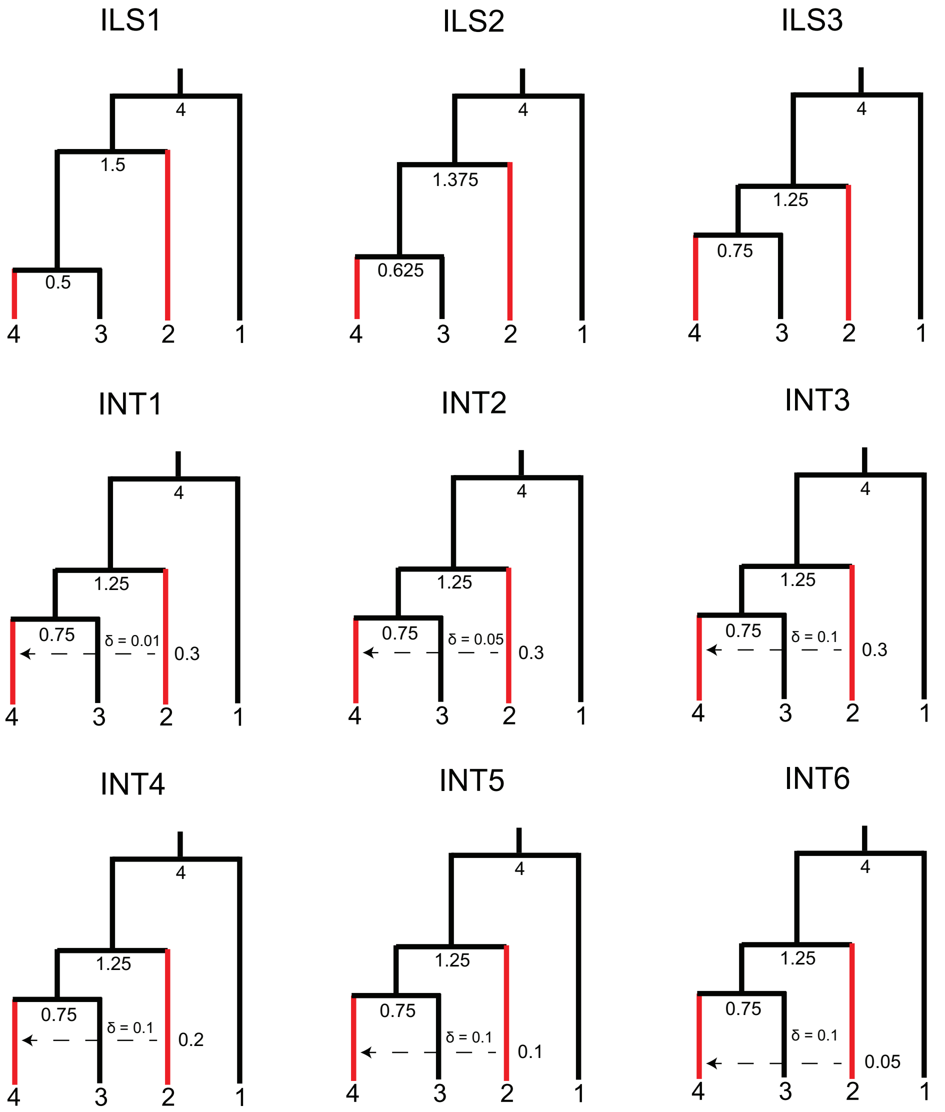


Supplementary Figure 2: Parameters used for benchmarking simulations in *HeIST*. Nodes and reticulations are labelled with the timing of the split in units of $2N$ generations. Branch lengths are visually adjusted in each condition to show how the parameters change, but they are not to scale. Species sharing the derived character are shown in red.

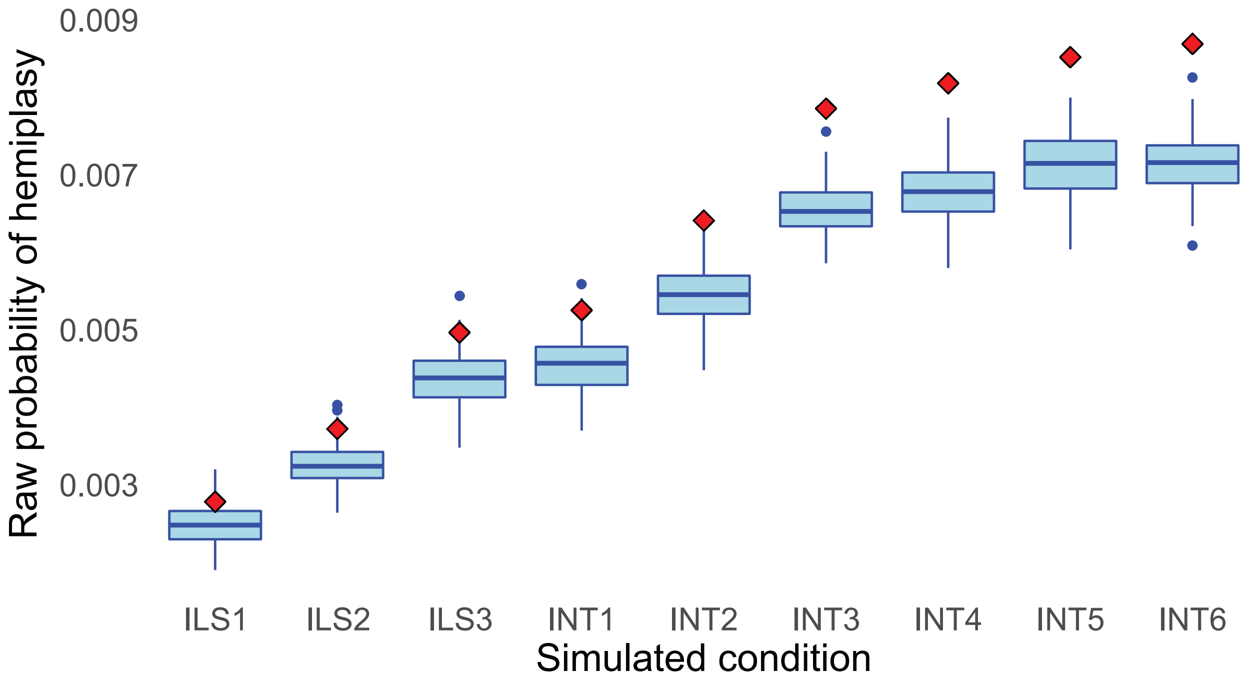


Supplementary Figure 3: Mismatch between simulated (blue boxes) and theoretical (red diamonds) raw probabilities of hemiplasy across simulated conditions.

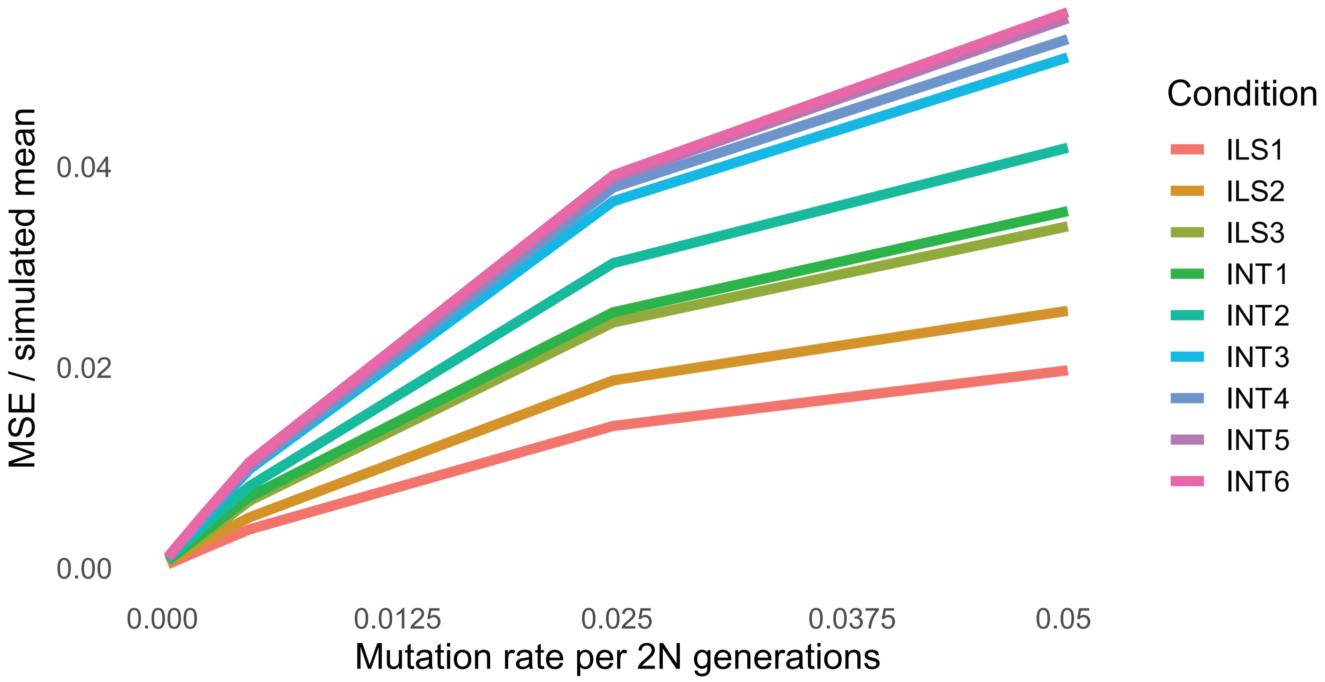


Supplementary Figure 4: Degree of mismatch between simulated and theoretical values of the raw probability of hemiplasy for our nine simulated conditions (colors) across five mutation rates (x-axis). In the y-axis, the mean-squared error between the simulated and theoretical value is normalized by the mean simulated value to make the error comparable across simulation conditions

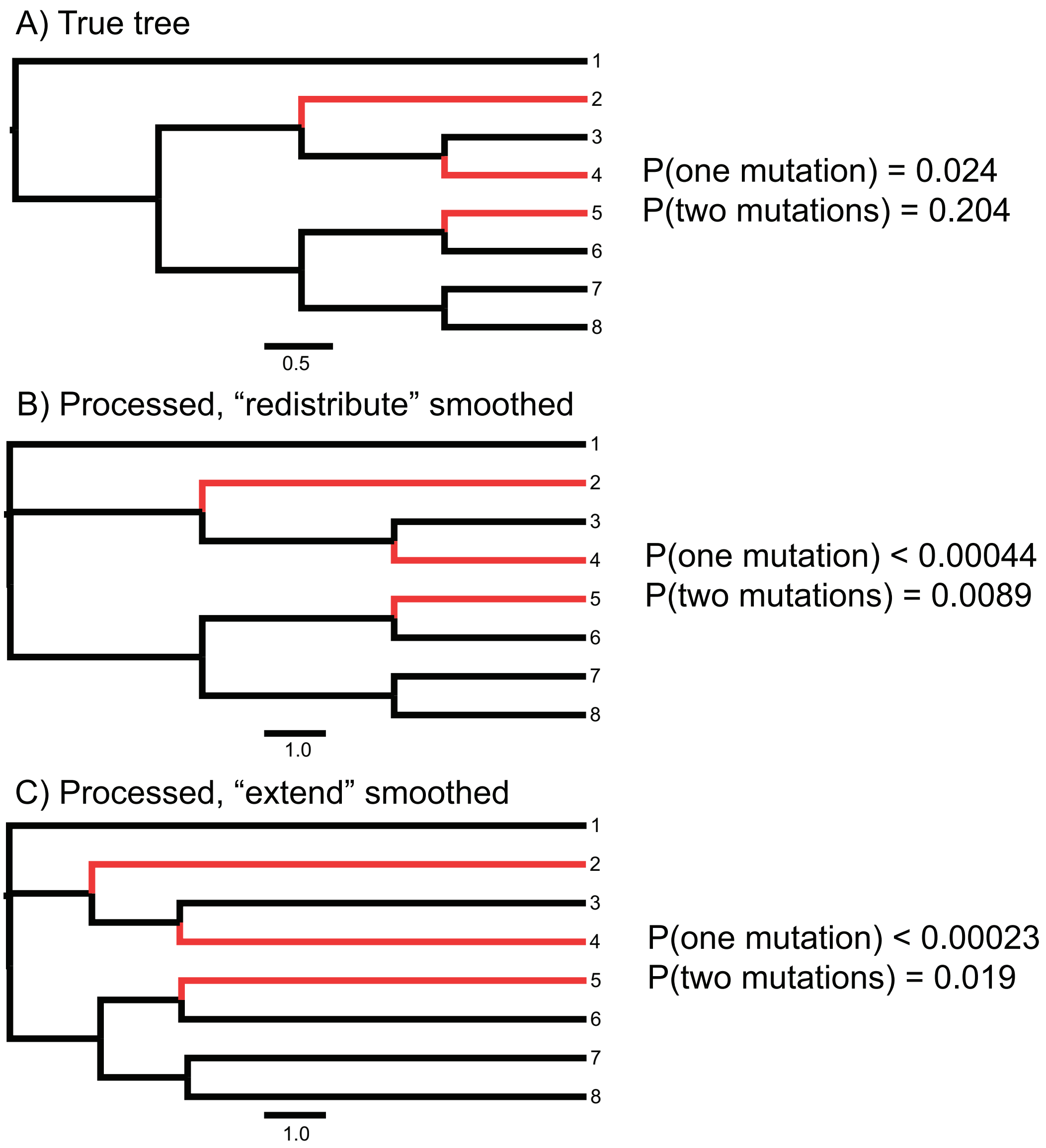


Supplementary Figure 5: Effect of phylogenetic inference, branch length unit conversion, and smoothing on estimated probabilities of hemiplasy in *HeIST*. Species sharing the derived state are highlighted in red. Panel A shows the true tree in units of $2N$ generations, with estimated probabilities for one and two mutations (i.e. hemiplasy cases). For panels B and C, gene trees were simulated from the tree in panel A and used to build a phylogeny with RAxML, which was then giving to the unit conversion module in *HeIST*. In panel B, smoothing was done by redistributing branch lengths, while in panel C it was done by extending the tip branches. Note the length scale is different in panels B and C than in panel A. The ancestral branch leading to the ingroup clade (all taxa except taxon 1) was originally inferred to be a polytomy in RAxML, but has been assigned a very short internal branch in the trees in panels B and C.

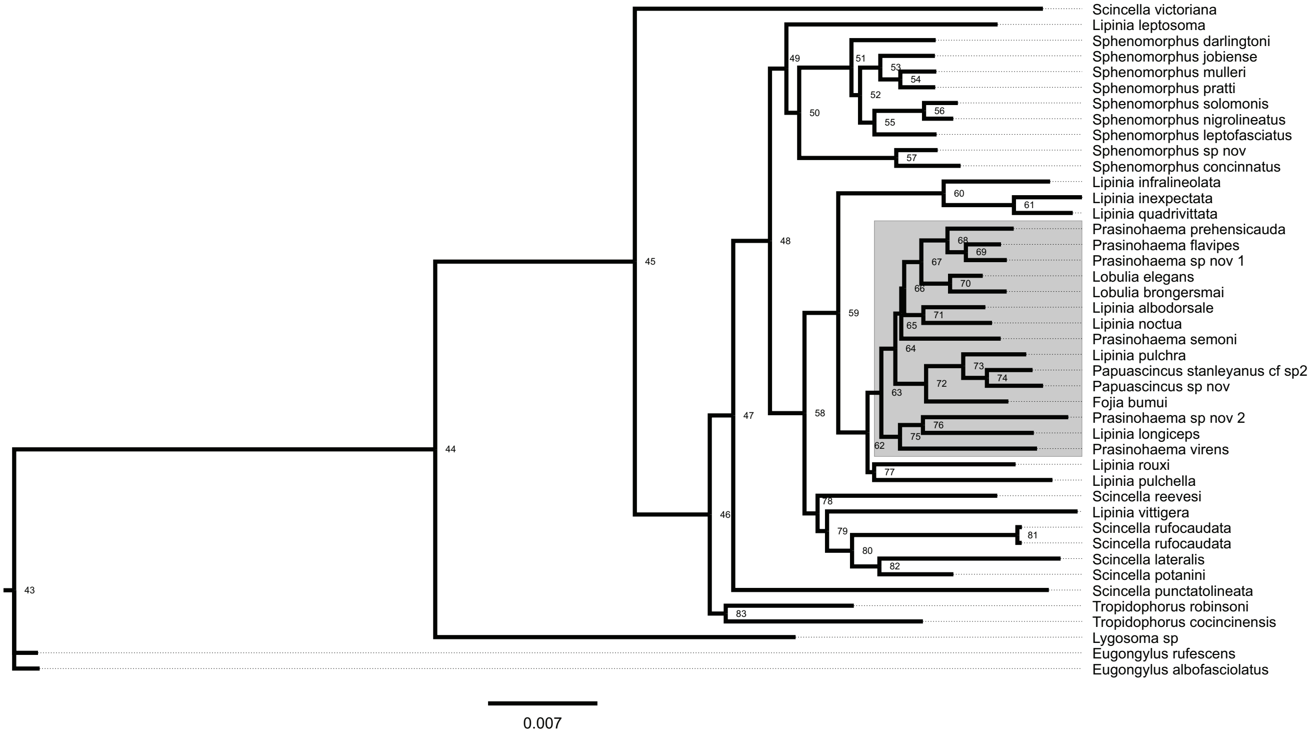


Supplementary Figure 6: Full 43-species maximum-likelihood lizard phylogeny constructed by RAxML, with branch lengths in units of substitutions per site. The 15-species subclade containing green-blooded species is highlighted in grey. Node labels correspond to the IDs in Supplementary Table 1 (below). This tree, and its associated site concordance factors in Supplementary Table 1, were used to obtain the linear formula for unit conversion in the regression module of *HeIST*.

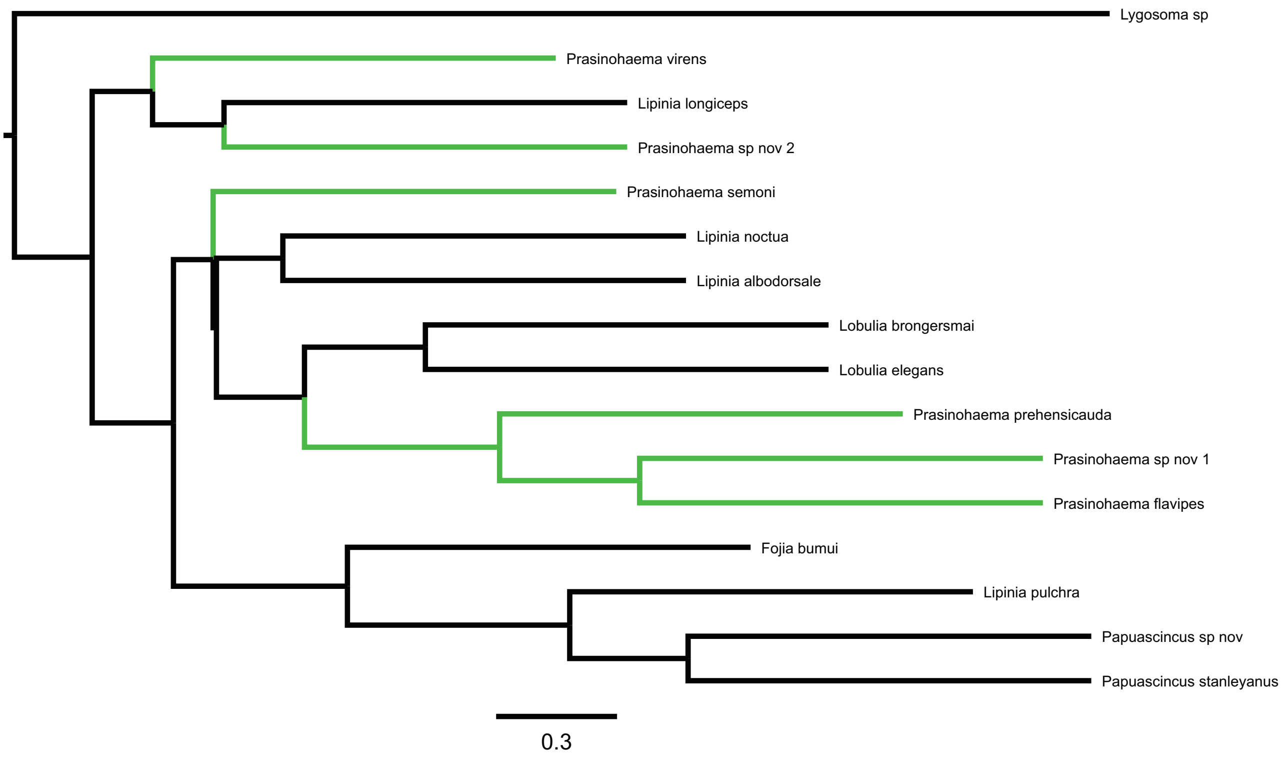


Supplementary Figure 7: Phylogeny of green-blooded lizards and an outgroup inferred from 3220 UCE gene trees using ASTRAL (data from Rodriguez et al. 2018). Internal branch lengths are shown in coalescent units of $2N$ generations, with arbitrary tip lengths of 1 coalescent unit. Taxa with green blood are labelled in green.

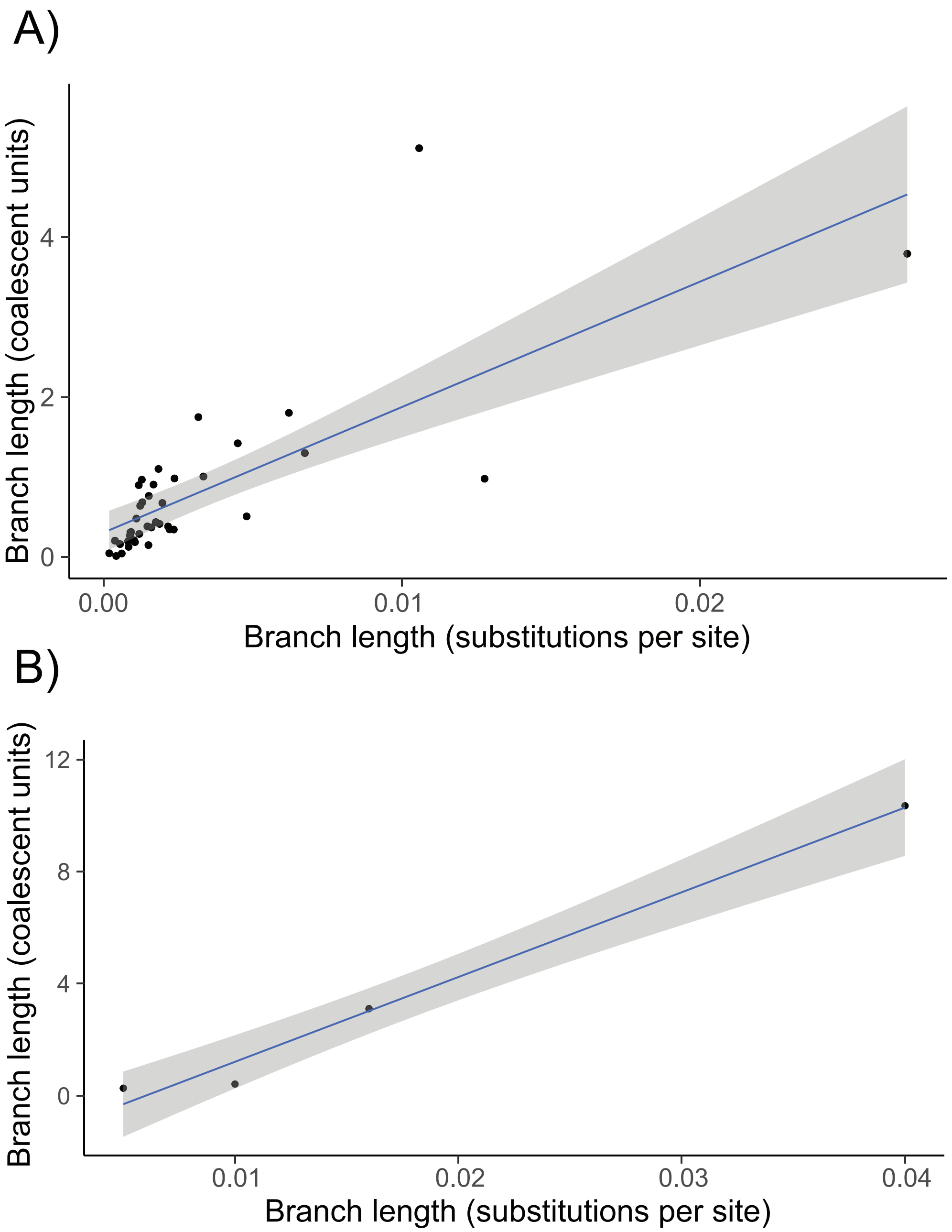


Supplementary Figure 8: Regression of internal branch lengths in substitutions per site (x-axis) against the same branch estimated in coalescent units using concordance factors (y-axis) for the 43-species lizard phylogeny (panel A) and the 6-species *Heliconius* phylogeny (panel B).

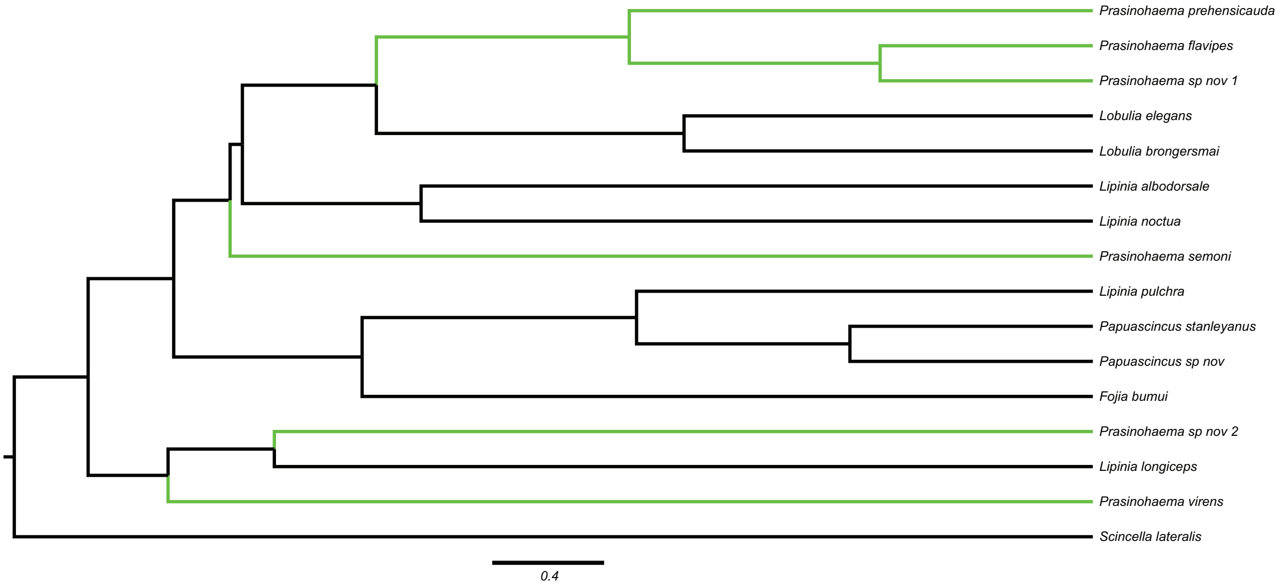


Supplementary Figure 9: Ultrametric phylogeny of green-blooded lizards, with branch lengths in units of $2N$ generations and green-blooded taxa labelled in green. This tree was generated from the phylogeny in Figure 1 in the main text using our regression approach for unit conversion, followed by a smoothing step. Gene trees were simulated from this tree within *HeIST*.

| ID | sCF | sDF1 | sDF2 | sN | Length |
|---|---|---|---|---|---|
| 44 | 98.51 | 0.83 | 0.66 | 13055.31 | 0.0269539 |
| 45 | 74.89 | 12.76 | 12.35 | 7471.64 | 0.0127777 |
| 46 | 59.84 | 20.29 | 19.87 | 2443.06 | 0.00479676 |
| 47 | 42.39 | 29.68 | 27.93 | 1758.14 | 0.00150258 |
| 48 | 52.59 | 23.33 | 24.08 | 2526.28 | 0.00235179 |
| 49 | 44.52 | 28.22 | 27.27 | 1792.81 | 0.00105067 |
| 50 | 44.96 | 26.38 | 28.67 | 1591.1 | 0.000819417 |
| 51 | 75.63 | 12.38 | 11.99 | 2027.75 | 0.0033454 |
| 52 | 43.07 | 29.26 | 27.67 | 877.28 | 0.000551229 |
| 53 | 66.33 | 16.75 | 16.92 | 1011.3 | 0.00129594 |
| 54 | 74.58 | 12.57 | 12.85 | 734.04 | 0.00128157 |
| 55 | 51.14 | 22.46 | 26.4 | 990.89 | 0.000919157 |
| 56 | 88.41 | 5.97 | 5.62 | 1255.3 | 0.00317694 |
| 57 | 88.96 | 6.12 | 4.93 | 3265.93 | 0.00621371 |
| 58 | 52.67 | 23.94 | 23.39 | 2055.31 | 0.00220984 |
| 59 | 54.37 | 23.38 | 22.25 | 1338.73 | 0.00216061 |
| 60 | 81.77 | 8.95 | 9.27 | 1655.73 | 0.00674802 |
| 61 | 83.93 | 4.85 | 11.21 | 576.59 | 0.00449816 |
| 62 | 55.76 | 21.13 | 23.11 | 1312.03 | 0.00187587 |
| 63 | 48.8 | 25.96 | 25.24 | 1213.47 | 0.000882666 |
| 64 | 50.85 | 24.71 | 24.44 | 1115.74 | 0.00089597 |
| 65 | 45.51 | 27.35 | 27.14 | 979.9 | 0.000379724 |
| 66 | 36.18 | 32.15 | 31.67 | 703.44 | 0.000186743 |
| 67 | 58.68 | 20.79 | 20.53 | 687.43 | 0.00109844 |
| 68 | 72.95 | 12.77 | 14.28 | 965.13 | 0.00167311 |
| 69 | 72.77 | 14.43 | 12.8 | 848.7 | 0.00117793 |
| 70 | 77.8 | 13.69 | 8.51 | 798.19 | 0.0018396 |
| 71 | 64.76 | 17.91 | 17.33 | 607.82 | 0.00122739 |
| 72 | 66 | 15.62 | 18.38 | 1489.12 | 0.00196959 |
| 73 | 74.96 | 12.1 | 12.94 | 1714.52 | 0.00237658 |
| 74 | 68.91 | 18.7 | 12.4 | 1212.9 | 0.00151573 |
| 75 | 49.94 | 24.67 | 25.39 | 1086.42 | 0.00119492 |
| 76 | 54.42 | 23.86 | 21.73 | 890.2 | 0.00146845 |
| 77 | 33.25 | 35.14 | 31.6 | 1213.04 | 0.000422247 |
| 78 | 41.07 | 27.48 | 31.45 | 1638.69 | 0.000836614 |
| 79 | 35.98 | 29.11 | 34.91 | 1548.07 | 0.000607296 |
| 80 | 53.75 | 23.01 | 23.24 | 1293.09 | 0.0016005 |
| 81 | 99.62 | 0.16 | 0.22 | 3067.45 | 0.0105833 |
| 82 | 56.8 | 19.86 | 23.33 | 724.33 | 0.00174971 |
| 83 | 45.68 | 27.14 | 27.18 | 772.84 | 0.00102174 |

Supplementary Table 1: Site concordance factors from IQtree. "ID": the ID of the internal branch in the full lizard phylogeny, as labelled in Supplementary Figure 5. "sCF": the value of the site concordance factor for the branch, averaged over 100 randomly sampled quartets. "sDF1" and "sDF2": the site discordance factors for the first and second most common discordant site patterns at each branch, respectively. "sN": the average number of informative sites averaged across the sampled quartets at each branch. "Length" is the length of the internal branch in substitutions per site.

| Trio | Species |
|---|---|
| 1 | Lipinia_longiceps.Prasinohaema_sp_nov_CCA01623.Prasinohaema_virens |
| 2 | Lipinia_longiceps.Prasinohaema_sp_nov_CCA01623.Prasinohaema_flavipes |
| 3 | Lipinia_longiceps.Prasinohaema_sp_nov_CCA01623.Prasinohaema_prehensicauda |
| 4 | Lipinia_longiceps.Prasinohaema_sp_nov_CCA01623.Prasinohaema_semoni |
| 5 | Lipinia_longiceps.Prasinohaema_sp_nov_CCA01623.Prasinohaema_sp_nov_CCA01005 |
| 6 | Lipinia_longiceps.Prasinohaema_virens.Prasinohaema_semoni |
| 7 | Lipinia_longiceps.Prasinohaema_virens.Prasinohaema_prehensicauda |
| 8 | Lipinia_longiceps.Prasinohaema_virens.Prasinohaema_sp_nov_CCA01005 |
| 9 | Lipinia_longiceps.Prasinohaema_virens.Prasinohaema_flavipes |
| 10 | Prasinohaema_sp_nov_CCA01005.Prasinohaema_flavipes.Prasinohaema_prehensicauda |
| 11 | Prasinohaema_sp_nov_CCA01005.Prasinohaema_flavipes.Prasinohaema_semoni |
| 12 | Prasinohaema_sp_nov_CCA01005.Prasinohaema_prehensicauda.Prasinohaema_semoni |

Supplementary Table 2: Trios involving green-blooded species which were evaluated for evidence of introgression using *D* statistics. Species are listed in the order P1, P2, P3, where an excess of shared P1/P3 or P2/P3 alleles would indicate evidence of introgression. *Lygosoma sp* was used as the outgroup for each trio.

| Trio | ABBA | BABA | D | stdev | CI-lower | CI-upper | p-val |
|---|---|---|---|---|---|---|---|
| 1 | 301 | 322 | -0.033707865 | 0.042090899 | -0.116206028 | 0.048790297 | 0.214 |
| 2 | 240 | 280 | -0.076923077 | 0.049594331 | -0.174127965 | 0.020281811 | 0.079 |
| 3 | 306 | 308 | -0.003257329 | 0.044616705 | -0.09070607 | 0.084191412 | 0.491 |
| 4 | 282 | 313 | -0.05210084 | 0.044610895 | -0.139538195 | 0.035336514 | 0.137 |
| 5 | 289 | 326 | -0.060162602 | 0.041842275 | -0.142173461 | 0.021848258 | 0.074 |
| 6 | 396 | 402 | -0.007518797 | 0.037634626 | -0.081282664 | 0.06624507 | 0.425 |
| 7 | 442 | 430 | 0.013761468 | 0.037376176 | -0.059495837 | 0.087018773 | 0.333 |
| 8 | 437 | 439 | -0.002283105 | 0.038342818 | -0.077435029 | 0.072868819 | 0.493 |
| 9 | 374 | 360 | 0.019073569 | 0.040634935 | -0.060570903 | 0.098718042 | 0.306 |
| 10 | 213 | 186 | 0.067669173 | 0.055838894 | -0.04177506 | 0.177113406 | 0.123 |
| 11 | 185 | 176 | 0.024930748 | 0.056183599 | -0.085189106 | 0.135050602 | 0.327 |
| 12 | 313 | 295 | 0.029605263 | 0.042237617 | -0.053180467 | 0.112390993 | 0.265 |

Supplementary Table 3: *D* statistic results for the 12 trios listed in Table 1, estimated from the concatenated alignment of ultra-conserved elements (UCEs). Significance was evaluated for each trio by bootstrap-sampling the UCEs to generate a null distribution of alignments, and asking how often the bootstrap distribution of 1000 *D* statistics was at least as extreme as the observed value.
